# An improved method for sampling and quantitative protein analytics of cerebrospinal fluid of single mice

**DOI:** 10.1101/2024.06.18.599559

**Authors:** Athanasios Lourbopoulos, Stephan A. Müller, Georg Jocher, Manfred Wick, Nikolaus Plesnila, Stefan F. Lichtenthaler

**Author notes:** equal contribution (Athanasios Lourbopoulos, Stephan A. Müller). **Sources of support:** Listed in the acknowledgment.

## Abstract

Mice are the most commonly used preclinical animal model, but protein analytics of murine cerebrospinal fluid (CSF) remains challenging because of low CSF volume (often <10 µl) and frequent blood contaminations. We developed an improved CSF sampling method that allows routine collection of increased volumes (20-30 µl) of pure CSF from individual mice, enabling multiple protein analytical assays from a single sample. Based on cell counts and hemoglobin ELISAs, we provide an easy quality control workflow for obtaining cell- and blood-free murine CSF. Through mass spectrometry-based proteomics using an absolutely quantified external standard, we estimated concentrations for hundreds of mouse CSF proteins. While repeated CSF sampling from the same mouse was possible, it induced CSF proteome changes. Applying the improved method, we found that the mouse CSF proteome remains largely stable over time in wild-type mice, but that amyloid pathology in the 5xFAD mouse model of Alzheimer’s disease massively changes the CSF proteome. Neurofilament light chain and TREM2, markers of neurodegeneration and activated microglia, respectively, were strongly upregulated and validated using immunoassays. In conclusion, our refined murine CSF collection method overcomes previous limitations, allowing multiple quantitative protein analyses for applications in biomedicine.

## Introduction

Cerebrospinal fluid (CSF) is the only body fluid in direct contact with the brain that is routinely accessible in a clinical setting. Alterations in CSF composition can inform about physiological and pathophysiological changes occurring in the brain. Consequently, CSF analytics for proteins or cells have become essential for diagnosis, prognosis, and treatment control of multiple neurological, neurodegenerative and psychiatric diseases and for a mechanistic understanding of the underlying pathophysiology^1^. Examples are the measurement of amyloid β (Aβ), total- and phosphorylated-tau for the diagnosis of AD^2^, and neurofilament light chain (NfL) as a marker for neurodegeneration ^3^. Human CSF is continuously produced at a rate of 300-600 μl/min and has a total volume of 90-150 ml that is turned over approximately 4-6 times per day^4^. Human CSF is easily accessible at larger quantities (at least 2-3 ml) in the clinical routine setting, which allows measurement of multiple analytes in parallel from the same collected CSF sample. Human CSF is collected via a needle that directly reaches the subdural space of CSF flow without contact to epidural areas that may potentially contaminate a CSF sample ^5^.

Mice are the most commonly used preclinical animal model. CSF protein analytics of single mice is possible, including whole proteome analytics using mass spectrometry (MS) ^6^, and may enable new biomarker discovery and validation, elucidation of mechanisms of disease pathogenesis and rapid translation of preclinical results to patients. Despite these promises, there are major challenges to mouse CSF protein analytics. Compared to humans, mice are smaller (20-40 g body weight), have a lower CSF production (0.32-0.35 μl/min)^7^, and less total CSF volume of 40 µl which is turned over approximately 12-13 times per day^8, 9^. Murine CSF collection requires special microsurgical techniques that fall into two approaches. One approach pierces the dura and allows open-CSF-flow on the epidural tissues before collection with a capillary (“epidural” or “open" collection)^10, 11^. The other approach comprises a clean dura-piercing and direct subdural collection of the CSF with a capillary with no extra-dural contact and contamination (“subdural” or “closed” collection) ^12^. This latter method yields mainly low volumes of CSF (mostly below 10 µl^13, 14^) and the former relatively larger volumes to a maximum of 10-20 μl^7, 13, 15^ from single mice. Yet, both methods are often below the sample volume requirements of ELISA-based assays. Thus, CSF samples are often pooled from different mice to obtain sufficient sample volume for protein analytics, which requires larger animal numbers and prevents the generation of single animal-resolved data^16^. Moreover, the available murine CSF sampling techniques (especially the “epidural” one) are prone to extra-CSF contamination, such as tissue proteins and blood, which prevents accurate measurements of many proteins in CSF because of the more than 150-fold higher plasma protein concentration (approximately 6.19±0.05 g/dl total protein) compared to CSF (approximately 26±1.5 mg/dl) in mice ^17, 18^ similar to humans (plasma: 6-7g/dl; CSF: <45 mg/dl)^19,20^. About 50% of CSF protein quantifications are affected by even low blood contaminations^21^. For human CSF, below 0.01% blood contamination is acceptable for global quantitative proteomics^21,22^. Low blood contaminations may not stain the CSF, making accurate visual recognition impossible and could lead to inaccurate results of CSF protein analytics^22^. Currently, no standard procedures are available to monitor blood contaminations of murine CSF.

To overcome limitations in murine CSF analytics, we developed an improved “subdural” (closed), contamination-free sampling method that now allows routine collection of 20-30 µl of pure CSF from individual mice and also enables repeated CSF sampling from the same animal. We provide an easy quality control workflow to ensure that mouse CSF is cell- and blood-free. This new method was applied a) to provide an estimation of absolute concentrations for hundreds of mouse CSF proteins using an absolutely quantified protein standard, b) to perform repeated sampling of individual mice to reveal resulting changes onto the CSF proteome and c) to identify CSF proteome changes induced by Aβ plaque formation using the 5xFAD mouse model of Alzheimer’s disease including further protein analytics with ELISA, and Simoa assays.

Collectively, this improved murine CSF collection method overcomes limitations of previous protocols^11, 12, 13, 15^, can be controlled to be free of protein or cellular contaminants for mass spectrometry-based biomarker studies, and allows multiple quantitative protein CSF analyses in parallel for many applications in neuroscience.

## Results

### Characterization and quality control of the collected mouse CSF

We first improved the procedure of blood-free, “subdural” CSF collection in C57BL/6 mice to increase the obtainable volume without CSF contact to the potentially contaminating epidural tissues. Higher CSF volumes (10-20 μl) have been reported only for the epidural collection methods^10, 11^. For the collection procedure, we compiled an operative set-up (Fig. 1A) including a classical stereotactic frame (Fig. 1A) to fix the mouse head in an optimal position (Fig. 1B) and a dissecting microscope to reveal the dura of the cisterna magna for CSF collection (Fig 1C). Critical points for collection of clean CSF are the fixation of the head, the careful exposure and cleaning of the dura in an atraumatic way without micro-bleedings, the tip of the capillary and the suction process of the CSF. The head of the mouse has to be bended at approximately 135° and be stable; the glass tip has to be broken to create a sharp and short bevel (Fig 1D) to pierce the dura mater of the cisterna magna but not touch the underlying brainstem; the suction force in the capillary has to be stepwise, minimal and smooth every approximately 5 minutes only when the collected CSF-column in the capillary has stopped increasing, if CSF pulsates and the cisterna magna is partially refilled. Eventually, the whole process lasts approximately 25-45 minutes for each animal. The collected volume depends primarily on the collection time (the longer the time, the larger the volume) and secondarily on the size of the cisterna magna of the animal. Using this method, we obtained 19-28 µl (median: 26) of clean CSF from adult wild-type C57BL/6 mice at different ages (3, 6, 12 months) (Fig. 2A). This is a significant improvement compared to the previous limit of 10-15 μl CSF per mouse reported in the few studies collecting CSF with the “subdural” method ^13, 15^, while most studies typically remain below 10 µl in total ^13, 14^.

**Figure 1.**
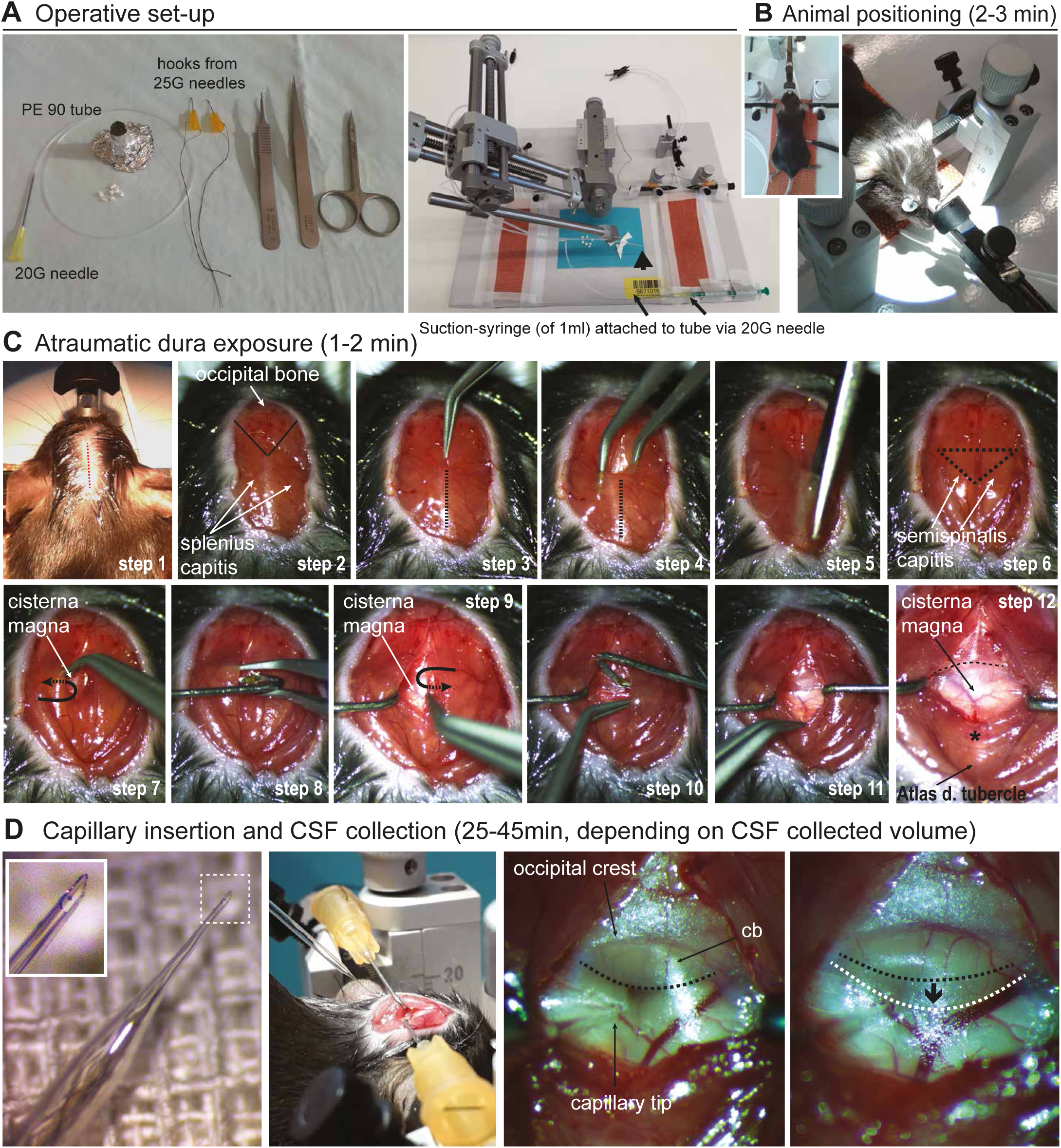
Stepwise and detailed experimental procedure of CSF collection with the subdural “closed” method . A) operative set-up required for cisterna magna puncture and CSF collection (see also text) and stereotactic device with the set-up of suction: 1 ml syringe, 20 G needle, PE90 tubing attached to the 20 G needle (syringe-side, arrows), the glass-capillary for dura puncture (arrowhead) and the pair of self-made orange hooks on ear-bars. B) The animal is placed on the ear- and nose-holder (insert in b), and is warmed by a temperature-controlled pad on the stereotaxic floor (the temperature probe is placed under the abdomen of the animal). Note that the head is bent at approximately 135°; whitish Bepanthen eye-cream is placed for cornea protection; the head is positioned under a stereotactic microscope. C) Atraumatic dura exposure (time required: 1-2 minutes) and detailed single steps (1 to 12, for description see text). D) Insertion of the glass capillary (first image and insert magnification: with the optimal shape of the sharp and fine glass-tip) in the cisterna magna (second and third image), followed by slow and stepwise CSF collection (fourth image, see text). Note the angular direction of the capillary towards the cerebropontine depth of the cisterna, that avoids any vessel (second and third images of D); collecting of CSF under mild suction results in mild caudal protrusion of the cerebellum (cb, white dashed line and arrow in fourth image of D) in comparison to its initial position (black dashed line in third image of D).

**Figure 2.**
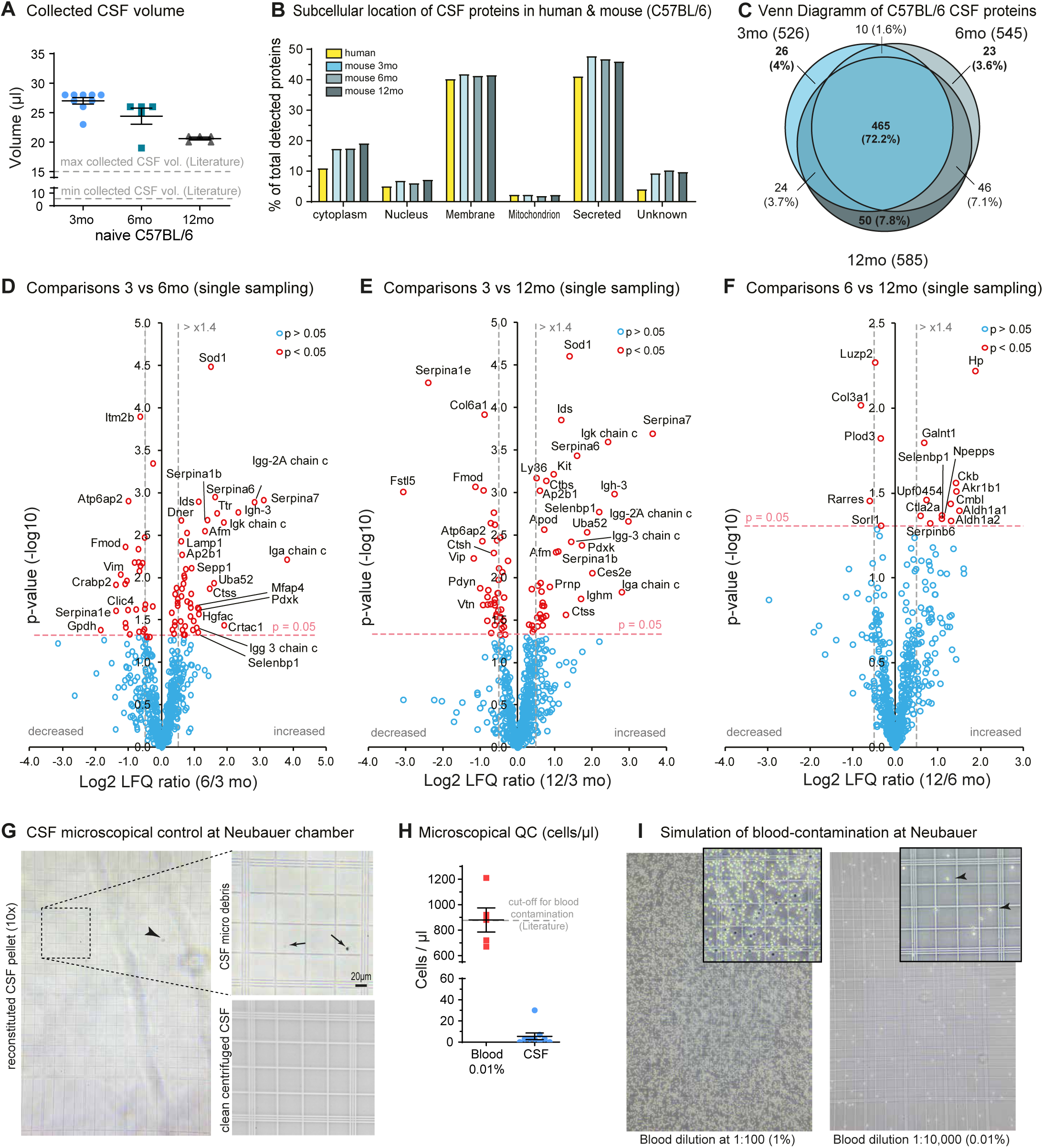
Quantitative and qualitative characterization of the mouse CSF. A) Collected CSF sample volumes (μl) for 3 (n=9), 6 (n=5) and 12 months (n=5) old C57BL/6 wild-type mice; dashed grey lines show the maximum and minimum reported collected CSF volume using the “subdural” contamination-free method. CSF was collected once from each animal (single-animal samples), no sample pooling was done. B) Qualitative comparison of mouse CSF to that of human, showing similar subcellular location of proteins in each group. C) Venn diagram of detected proteins in mouse CSF of different ages, showing that 72.2% of them are detected through different ages of mice. D-F) Volcano plots of CSF-proteome comparisons (6 versus 3, 12 versus 3 and 12 versus 6 month-old C57BL/6 mice): x-axes show the Log2 change in protein LFQ relative intensities (increased or decreased abundance) and y-axes the minus log10 p-value (two-sided Student’s Ttest) for each single protein of the plot; red open-circles indicate significantly changed proteins (p<0.05), blue are not significant. Note that most of the changes occur between 3 and 6 months of age (D). G) Microscopical control of CSF’s pellet under a 10x and 20x (magnifications to the right) objective. This pellet contains the particles or cells that were in the collected CSF, before centrifugation. Note that apart from few debris (arrows and arrowheads), which is presumably due to dura puncture and is removed by centrifugation, the CSF is largely devoid of cells. H) Counting of CSF-cells (in its reconstituted pellet) showed presence of 0-3 cells (see text) per μl (n=9, mean ±95%CI) which is significantly lower than the 0.01% blood contamination cut-off criteria (n=5, mean ±95%CI). I) Simulation experiments of a blood-contamination with 1% and 0.01%, as shown in a Neubauer chamber (under a 10x objective and 20x insert magnification, arrowheads point at erythrocytes); cellular results for the 0.01% contamination are shown in (H).

Applying mass spectrometry-based proteomics, the majority of CSF proteins were annotated as either secreted soluble proteins or proteolytically released membrane protein ectodomains, in agreement with relevant previous studies, where less than 15 µl of mouse CSF were obtained using the “subdural” approach ^23, 24^ (Fig. 2B). A similar protein distribution was seen in human CSF (Fig. 2B). Whole proteome analysis revealed that the mouse CSF proteome remains largely stable between 3 and 12 months of age, with some changes occurring mainly between 3 and 6 months (Fig. 2C, and 2D-2F, Suppl. Data 1, Suppl. Fig. 1).

To assess the purity of the CSF, we counted how many cells it contains per microliter and how this number would change upon blood contamination, which may happen during CSF collection. CSF, freshly taken from the 3-month-old C57BL/6 mice used above (n=9), was centrifuged for 10 min at 2000 g and transferred to a new tube. The CSF was clean and had no red color. A pellet, which would indicate precipitated cells, was not obviously visible after centrifugation. Yet, we considered that mouse CSF may contain very few cells so that the resulting pellet would be too small for visual detection. To test this possibility (and keep the CSF for analyses), we resuspended the presumed pellet in PBS and counted indeed 1.1±1.2 cells per 25 small Neubauer squares (range 0-9), which is equal to 3.7±4 cells/μl in the original volume of collected CSF (Fig. 2G and 2H). The reconstituted pellet additionally contained isolated macro- or micro-debris (magnification in Fig. 2G, presumably dura-puncture related).

To analyze how a possible blood-contamination would affect the cell count in CSF samples, we carried out simulation experiments diluting mouse blood in PBS. A 1:10,000 blood dilution (0.01 %), that is acceptable for proteomics experiments of human CSF^22^, resulted in 16.6±4.5 cells per 5 small Neubauer squares, equal to 880±212 cells/μl (mainly RBCs) (Fig. 2H and 2I). This number of cells was much higher compared to the 3.7±4 cells/μl found in our collected CSF, as described above. Thus, the collected CSF contained at least 200-times fewer cells, compared to the currently accepted cut-off quality value of 0.01% for cellular contamination ^21, 22^. We conclude that these cells constitute the normal cellular component of murine CSF, similar to human standards, which report 5 cells/µl as normal^5^.

The cell-free CSF obtained after centrifugation may contain microparticles, such as extracellular vesicles, which was investigated with dynamic light scattering (DLS). Mouse CSF showed three constant peaks at 10.4±1.1, 73.9±21.1 and 687.8±178.8 nm before centrifugation, that were preserved after centrifugation at 800 g (11.1±0.9, 69.9±18.0 and 513.4±148.0 nm) and 2,000 g (9.0±0.9, 69.6±3.4 and 487.3±132.1 nm) (Suppl. Fig. 2A). These CSF peaks correspond to microparticles with an approximate size range of 10, 70 and 500 nm, that are much smaller than cells. Particle analysis of human CSF samples (Suppl. Fig. 2B) showed a similar microparticle distribution after centrifugation at 800 g (10.3±0.9, 67.3±26.4 and 343.8±20.0 nm) and 2000 g (10.9±0.6, 65.6±16.6 and 385.1±36.3 nm), in accordance with data reported in the literature in the range of 26-305 nm for human CSF, depending on the measurement method and underlying CSF condition^25^.

Collectively, our results indicate that our method surpasses previous ^13, 15, 22^ quantity and quality standards of CSF murine collection. Quality control for possible blood-contamination using the classical Neubauer chamber in the CSF “pellet” is fast, reliable, sensitive, does not use valuable CSF sample and is translational to human standards^5^.

#### Repeated mouse CSF collection from single mice

Next, we tested if repeated CSF sampling of the same mice is possible, and whether this procedure affects the CSF proteome (see experimental design in Figure 3A). We collected CSF from 3-month-old wild-type mice and from the same mice again at 6 months. At 3 months, 25.8±1.7 μl of CSF volume were obtained (Fig. 3B), in line with the results from singly punctured mice (Fig. 2A). Upon repeated sampling at six months, we collected 19.0±0.9 μl of clean CSF. Sampling at intervals shorter than three months after the initial sampling did not appear reasonable, because the dura strongly reacted to the initial puncture with fibrosis and thickening and complementary shrinkage of the cisterna magna, which was evident during the first days after the initial puncture (Figure 3C, middle photo) and lasted for at least four weeks. After three months (at the age of six months), the dura and cisterna magna were macroscopically restored (Figure 3C, right photo) and CSF sampling was feasible, in accordance with a previous report^14^.

**Figure 3.**
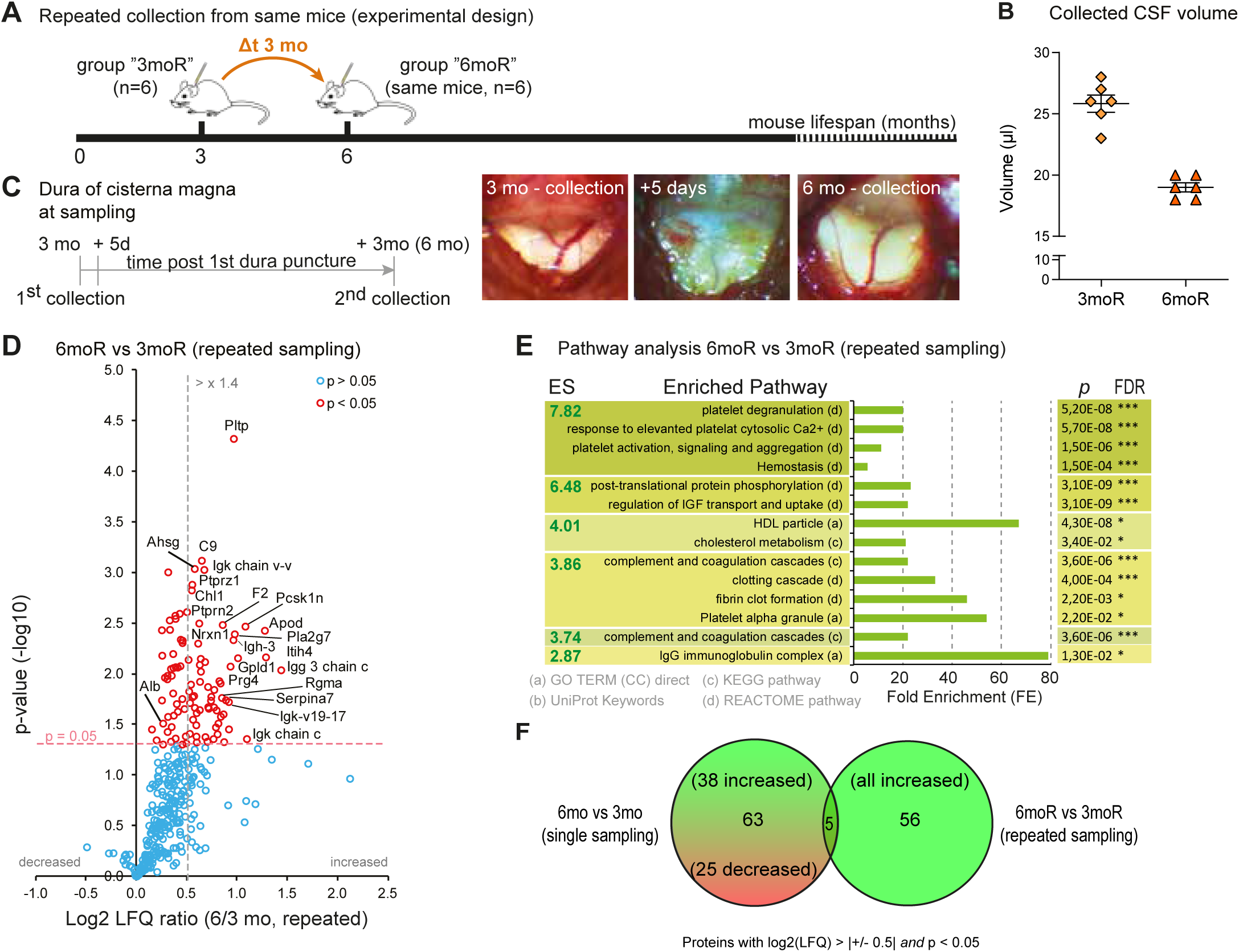
Repeated mouse CSF collection from single mice. A) CSF collection from C57BL/6 mice at 3 months (3moR) followed by a second collection in the same animals at 6 months (repeated collection, 6moR), "repeated collection" experiment (n=6). B) collected CSF volume at each timepoint (at 3 and then 6 months) from each mouse (mean ±95%CI). C) timeline and representative photos of dural appearance at first CSF collection (3 months, 1st collection) 5 days after the collection (photo of dura: note the intense fibrotic scarring and the "whitening" of the dura) and at the 2nd collection 3 months after the first (6 monthś old mice, the dura in now clear and cisterna magna fully reconstituted). D) Volcano-plot diagram of CSF proteomic changes between 6 (repeated collection, “6moR”) and 3 (“baseline” collection, “3moR”) months in the same animals (red open-circles indicate significantly changed proteins, one sample T-test with Ho=0 p<0.05); note an overall increase of proteins mass (positive Log2 LFQ ratios). E) Functional annotation clustering analysis by DAVID-Database for the repeated sampling cohort only, as this is shown in (D) (ES = Enrichment Score); green boxes indicate increased pathways (note that the majority of proteins are increased by the repeated collection); the right columns show the corresponding p- (Benajmini correction) and FDR-(false discovery rate) values for each enriched (i.e. changed) terms (*<0.05, **<0.01, ***<0.001). Despite a fully reconstituted cisterna magna, the CSF showed a significantly changed proteome. F) Venn diagram of the significantly altered proteins (proteins with significant log2 LFQ increase of more than ±0.5) in single and repeated sampling cohorts; the two cohorts were significantly different both in terms of changed proteins and the direction of change (increased or decreased).

The CSF proteome analysis (Suppl. Data 1) showed that the repeated sampling at six months increased the overall protein content compared to the 3-month time point, which is demonstrated by the right shift of the volcano plot for repeated sampling (Figure 3D) in comparison to the symmetric volcano of single sampling (Figure 2D). The increased protein content is likely a response to the fibrosis and thickening of the dura as seen macroscopically (Fig. 3C).

Next, we compared the proteome changes detected for repeated and single sampling cohorts, to understand if the repeated sampling at six months from the same mice (proteome changes in Figure 3D) changed the CSF differently compared to single sampling of different mice at six months (proteome changes in Figure 2D). Therefore, we analyzed only those proteins with a significantly altered abundance (log2 fold change > |+/- 0.5| and p < 0.05). We found that the two cohorts showed different proteomic signatures (Figure 3E, F): 63 proteins (38 up, 25 down) were uniquely altered for single CSF sampling, while 56 proteins uniquely altered (all increased) for repeated CSF sampling (Figure 3E, F). On the other hand, they had only a small overlap of five proteins in common (Figure 3F), four of which increased in both samplings (Igh-3, Ig kappa chain C region, Ig gamma-3 chain C region, and Serpina7) and only one changed in the opposite direction (Pcsk1n), being more abundant for repeated CSF sampling and less abundant for single sampling at the 6 versus 3-month time points. Conclusively, CSF sampling of individual mice induces proteome changes, which cannot be explained by aging from three to six months alone, but rather by the previous CSF collection at three months. This CSF-proteome change should be taken into consideration when CSF-proteome studies with repeated collection are planned.

### Relative and estimated quantification of mouse and human CSF proteins

The concentrations of very few murine CSF proteins are reported with absolute values, including total protein (adult, 27.7±0.6 mg/dl, 26±1.5 mg/dl)^17, 18^, Albumin (17.1±0.7 mg/dl)^17^, ApoE (approximately 1.2-6 μg/ml)^26, 27^, GFAP (approximately 100-200 ng/l)^16^, Apoa1 (approximately 0.60 μg/ml)^27^, transthyretin (35 μg/ml in APPswe/PS1A246E mice)^28^, decorin (approximately 10-30 ng/ml)^29^. Because of the lack of absolute concentrations, the majority of murine CSF studies refer to relative changes between different groups analyzed and prohibit comparisons between studies and laboratories. The lack of absolute concentrations is mainly due to the restricted volumes of collected CSF for classical biochemical assays, such as ELISA.

To estimate absolute concentrations of mouse CSF proteins, we set up a method that we validated with human CSF where larger CSF volumes are available. First, to provide an initial comparison of murine and human CSF proteomes we performed an analysis based on intensity-based absolute quantification (iBAQ) intensities, (Fig. 4A), a rough estimate for the protein abundances within a sample, taking the data obtained from mice at 3, 6, and 12 months (see Fig. 2) and eight human CSF samples. Selected iBAQ data are shown in Fig. 4B for APP and apolipoprotein E (ApoE), who are linked to cardiovascular and Alzheimer’s disease ^30^. A comparison of estimated CSF protein concentrations with reported values are shown in supplementary table 2.

**Figure 4.**
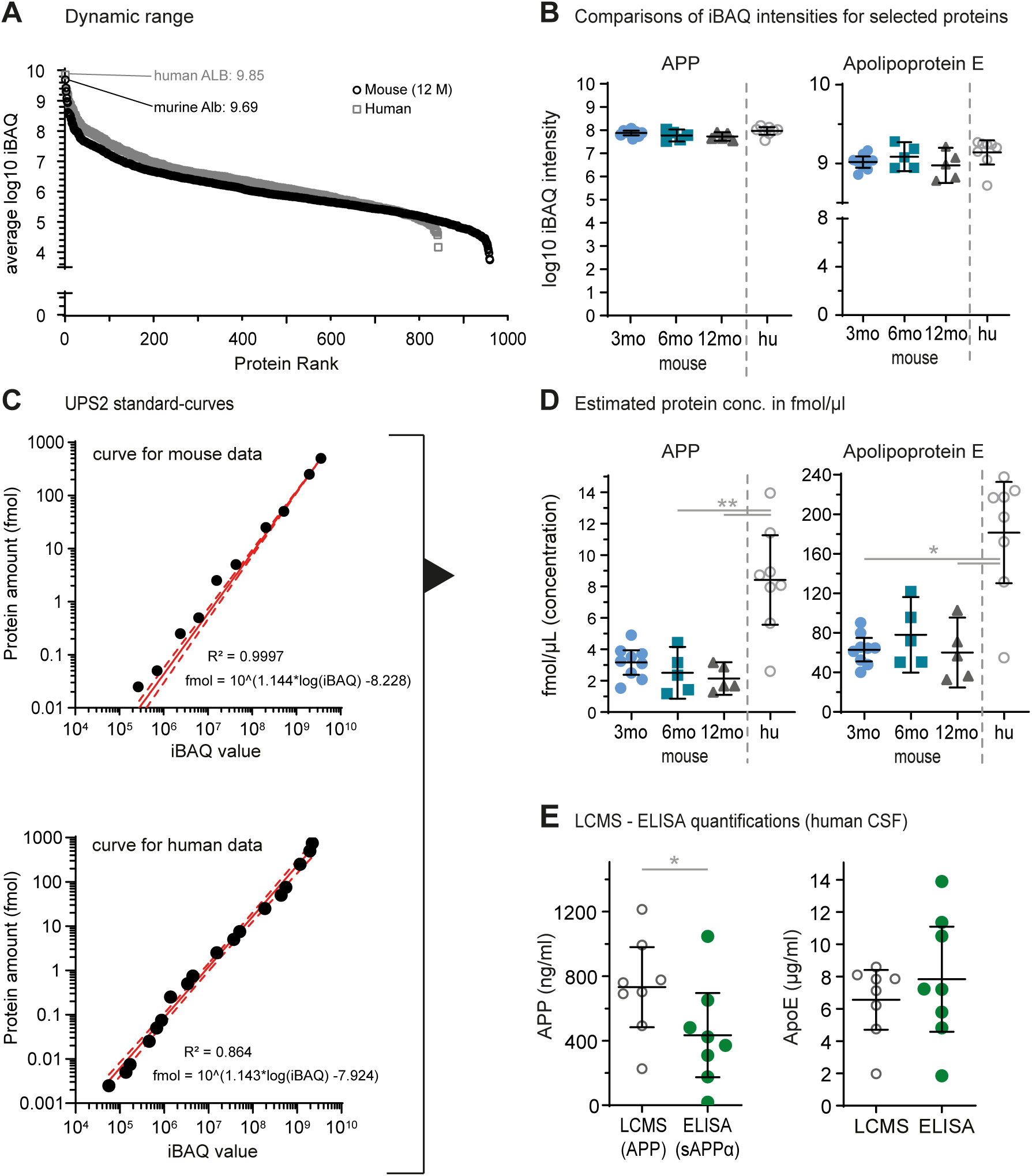
Relative and estimated quantification of mouse and human CSF proteins. A) Dynamic range of proteins for mouse (2 μl of sample, 12 months) and human (2 μl of sample) CSF, shown as log10 transformed iBAQ values indicate a detection range of about five orders of magnitude. B) Comparisons of quantified selected common or abundant proteins in CSF of 3 (n=9), 6 (n=5) and 12 (n=5) months old mice and humans (n=8) using log10 transformed iBAQ intensities. APP and apolipoprotein E are detected at higher levels in human CSF compared to mice. C) Plotting of standard iBAQ-fmol curves including the calibration equations for mouse and human sample analysis, based on values obtained from the proteins of the UPS2-kit. These equations were used to estimate the concentrations of proteins based on their iBAQ intensities. D) Estimated concentrations (fmol/μl) for the proteins shown in (B). E) Corresponding quantification of ApoE and sAPP in human CSF via classical ELISA assays show similar results to those of LC-MS. Graphs show mean ±95%CI of each group where applicable; * = p<0.05, ** = p<0.01, *** = p<0.001; ns = not significant.

To estimate absolute molar concentrations of all CSF proteins, we used the UPS2 kit proteomics dynamic range standard set, which covers 5 orders of magnitude with 48 human proteins, on the basis of their iBAQ intensities. Previously, the kit was applied for protein concentration quantification e.g. in mouse brain homogenates^31^, human cells^32^, E. coli^33^, HeLa cells^34^ or mouse stem cell^35^ samples, as well as validation of an Orbitrap benchtop mass spectrometer^36^. Based on these external standards measured together with the sample batches of murine or human CSF, we obtained calibration curves plotting the protein amounts against the iBAQ intensities with a high linear correlation (Fig. 4C). Based on the derived equations (see equations in Fig. 4C) we estimated concentrations (fmol/μl) for all detected mouse and human CSF proteins on the basis of their iBAQ values (Suppl. Data 2 and 3). According to the calibration range of the UPS2 standard, we could estimate the concentrations of 422 murine and 502 human CSF proteins. Representative human proteins (APP and ApoE) as well as selected mouse proteins are shown in Fig. 4D.

To validate the protein concentration estimation based on iBAQ intensities, we used human CSF, where larger volumes of CSF were available. ELISAs for human ApoE and soluble APPα (sAPPα) yielded concentrations comparable to the estimated concentrations (Fig. 4E), open circles estimated concentrations, green dots measured with ELISA) and were also in accordance to previous reports for ApoE (approximately 3-11 mg/l for adults^37, 38^) and total sAPP (approximately 750 ng/ml)^39^, respectively. Furthermore, an extended comparison of the estimated CSF protein concentrations (in fmol/μl) for selected human CSF proteins showed values similar or very close to those reported in literature (Suppl. Data 3). Some absolute quantification estimates of murine proteins in our study showed a concentration range similar to values reported in literature. This includes ApoE (estimated 1.6-2.7 μg/ml, reported approximately 1.2-6 μg/ml)^26, 27^ and GFAP (estimated 0.1-0.2 ng/ml, reported approximately 0.1-0.2 ng/ml)^16^. For transthyretin, a protein with a very high concentration close to the upper limit of the linear range, we obtained lower estimated concentrations (estimated 0.5-2.8 μg/ml, reported approximately 35 μg/ml in APPswe/PS1A246E mice)^28^. This indicates that iBAQ intensities provide a rough concentration estimation.

Nevertheless, although absolute quantification values are normally provided by targeted quantification based on absolutely quantified isotopically labeled peptides, the estimation using the UPS2 standard as a reference delivers values for hundreds of CSF proteins and might serve as an indicator of protein concentrations for further targeted assay development.

### Multiple protein analytical assays are feasible with single CSF samples

Finally, we applied our improved CSF sampling method to showcase that multiple types of protein analyses are possible with one collected sample of mouse CSF. Therefore, we sampled CSF of the 5xFAD^40^ transgenic mouse model of Alzheimer’s disease and compared its CSF proteome to the one obtained from corresponding wild-type littermates (wt) at seven months of age followed by validation of key findings using orthogonal methods (ELISA, Simoa) (Fig. 5A).

**Figure 5.**
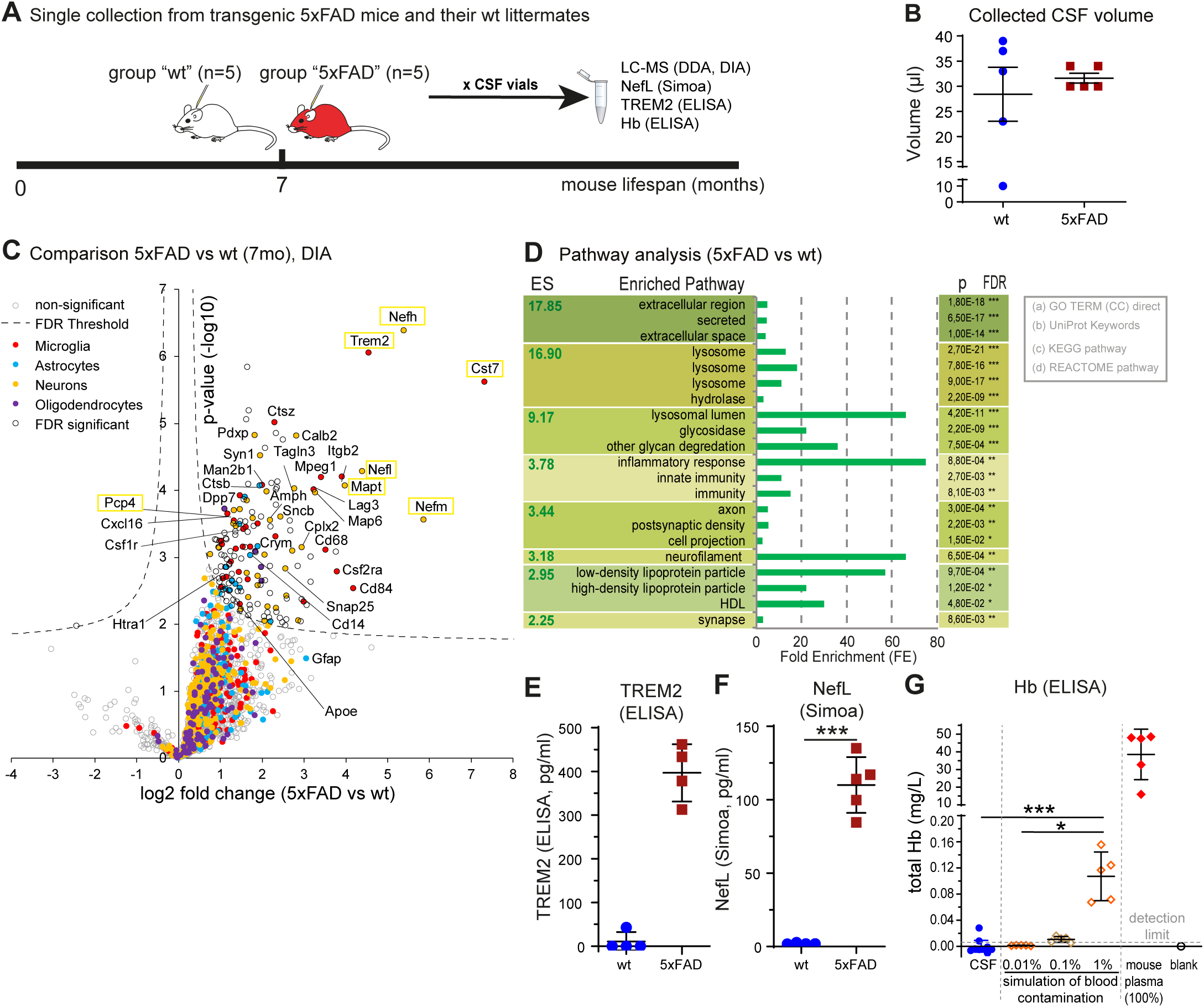
Multiple protein analytical assays are feasible with single CSF samples. A) single-sample CSF collection from wt and 5xFAD animals at 7 months of age (no pooling). CSF was aliquoted in multiple vials (5μl each) and 4 different assays were used (see text). B) collected CSF volume in each group (wt and 5xFAD) from each mouse (mean ±95%CI of each group; one wt animal fell for technical reasons below the techniqués minimum volume). C) Volcano-plot of CSF proteomic changes (x-axis: log2 protein LFQ change, y-axis: -log10 p-value, two-sided Student’s Ttest) between 5xFAD and wild-type using data-independent analysis . The dashed line indicates a permutation based FDR correction (p=0.05; s0=0.1). Proteins are categorized and labeled by color, based on their major cell type of origin^24^ . Selected and neurodegeneration-relevant proteins are highlighted in yellow boxes. Many show a strong increase, e.g. neurofilaments, Trem2 or MAPT (tau) in 5xFAD animals. D) Pathway analysis between 5xFAD and wt using the functional annotation clustering tool of DAVID-Database (same methodology and nomenclature as in Figure 3). E-F) Verification of selected quantified proteins, Trem2 (E), Nefl (F) with MSD ELISA and Simoa, respectively. G) ELISA-based quantification of hemoglobin (Hb) in CSF of both wt and 5xFAD animals shows levels below the detection limit of the method for 9/10 samples; for comparison our simulation experiments of blood contamination show detectable Hb for 1% contamination, and below detection limit for 0.01% (minimal accepted level of CSF contamination with blood^22^). Normal mouse plasma contained 38.5±14.3 mg/L of free Hb.

The sampled CSF volume was approximately 30 µl (Fig. 5B). For a whole proteome analysis, we used 5 µl, of which half of the digested sample was used for data-dependent acquisition (DDA) and data-independent acquisition (DIA) on a Q-Exactive HF mass spectrometer. Due to the pronounced pathology at seven months in 5xFAD mice, many proteins showed missing quantification values for wild-type control samples, likely because they were below the detection limit, but complete profiles for 5xFAD mice. Therefore, we filtered for proteins with complete quantification data in either the wt or 5xFAD group and imputed missing values with the software Perseus applying a downshift of 1.8 based on a normal distribution (Suppl. Data 4). Finally, 1696 protein groups were relatively quantified after data filtering and imputation. The volcano plot indicates increased protein content in the 5xFAD CSF with numerous proteins showing significantly increased abundance (Fig. 5C). Albumin levels were similar between wt and 5xFAD mice (Suppl Fig. 3C). The increase of several microglial proteins such as Trem2, Cst7, CD84, Lag3, Lyz2, Ctsz, Ctsa, and Ly86 is indicative of a strong microglial activation. These changes are in line with the fact that 5xFAD mice at seven months show plaque pathology, microgliosis, cognitive impairment, and synaptic losses^40, 41^, and similar to those obtained in another AD mouse line, APPPS1, at 12 months of age^42^. Additionally, the neuronal neurofilament proteins L, M and H (Nefl, Nefm and Nefh) and tau (MAPT), proposed markers for neurodegeneration, as well as the Purkinje cell protein 4 (PCP4) showed an increased abundance (yellow boxes Fig. 5C; for selected proteins see also Suppl. Figure 3). A pathway and gene ontology enrichment analysis showed that the amyloid pathology in 5xFAD mice at 7 months of age had upregulated pathways related to extracellular space and secreted proteins, lysosome, glycosidase, glycan degradation, inflammatory and innate immune responses, axons, neurofilaments, synapses and postsynaptic density, as well as lipoprotein particles (Figure 5D). Collectively, these changes suggest a starting damage of neurons and neurodegeneration already at 7 months of age, even though neuronal loss was previously described at a later time point ^43^.

Since our new CSF collection method provided enough CSF volume per mouse for further protein analytics, we validated two protein changes in the same CSF samples by orthogonal methods. We chose Nefl and TREM2, which are under evaluation as potential key markers for neurodegeneration and microglial activation in Alzheimer’s disease^10, 44^. Nefl was detected by SIMOA immunoassay using 3.4 µl of single mouse CSF. TREM2 concentration was measured by ELISA in 10 µl of single CSF samples. Similar to the proteomic results (Fig. 5C), we found significantly increased levels of Nefl and TREM2 in the CSF of 5xFAD mice (Fig. 5E, F), as a result of neurodegeneration and inflammatory microglial activation. The detected increases were in accordance with their corresponding relative quantification by nLC-Mass Spectrometry (Suppl. Fig. 3D, F).

Using 1 µl of the remaining mouse CSF, we also measured hemoglobin (Hb) by ELISA to determine whether an Hb ELISA may be useful as an alternative to the cell counts of CSF in the Neubauer chamber (Fig. 2) to ensure that CSF was free of a blood-contamination. Low levels of Hb are expressed endogenously in brain and may be found in CSF^45^, but are expected to increase dramatically upon a blood contamination. The Hb ELISA values from 1 µL of CSF were below the detection limit equal to the “blank values” (Fig. 5G). As a control, we simulated how a blood contamination would enhance the CSF concentration of Hb. We artificially diluted mouse blood into PBS at a dilution of 1:100 (1%) and 1:10,000 (0.01%) and compared the ELISA Hb measurement to pure mouse plasma. While the ELISA assay detected Hb in plasma (38.5±14.3mg/l) and at 1% contamination (0.11±0.04mg/l), the Hb values were below the method’s detection limit for the artificial 0.01% contamination ^21, 22^ (Fig. 5H). The data indicate that ELISA-based Hb measurement is able to monitor a potential blood contamination of CSF that may affect CSF proteomics.

Our data suggest that mouse CSF can be collected in a blood-free manner and that the ELISA-based Hb measurement can be taken as an alternative to cell counting for controlling the quality of collected mouse CSF. Moreover, the increased CSF volume obtained with our method allows multiple protein analytics assays to be performed with single samples. Therefore, we propose a sample collection scheme using centrifugation of CSF and a quality control of the potential reconstituted cell pellet with a Neubauer chamber as a gold standard (Fig. 6). Alternatively, a hemoglobin ELISA may be used as quality control, for example for samples that have previously been collected and where the Neubauer chamber analysis is no longer possible (Fig. 6).

**Figure 6.**
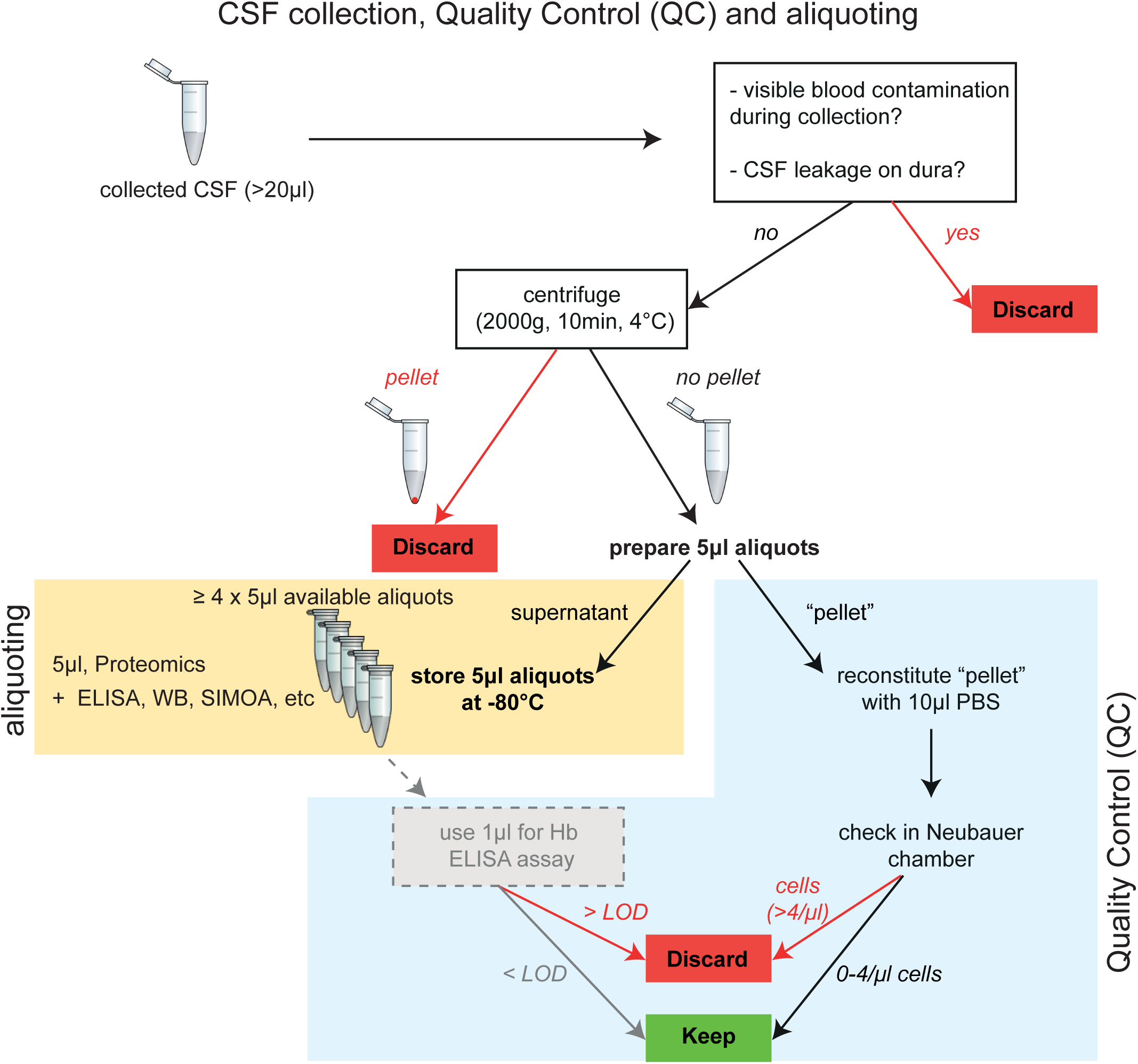
Quality control scheme for collection and handling of murine CSF samples. CSF is aliquoted in 5 μl aliquots in multiple low-protein-bind Eppendorf tubes, for subsequent use for mass spectrometry, Simoa, ELISA, and at least 1-2 more molecular techniques requiring 5μl of sample volume each. If a visible pellet is evident after centrifugation, samples are contaminated with cells and should be discarded. For quality control, we suggest investigating the cellular content of its reconstituted pellet (after centrifugation) with a Neubauer chamber, which does not consume CSF. If more than 4 cells/μl are detected, the sample should not be used for mass spectrometry analysis. A post-aliquoting controlling of CSF quality (i.e. blood contamination during the collection procedure) can also be done with Hb-ELISA: the detected Hb levels should be equal to blank (i.e. below level of detection, LOD).

## Discussion

CSF analytics provides essential information about the status of the healthy and diseased brain. Clinical CSF analysis in humans is widely used to diagnose neurological, psychiatric, and neurodegenerative diseases. Mice are the most common preclinical animal model. Murine CSF analytics has been used to study mouse models of brain diseases and the consequences of gene deletions ^6, 23^ ^24, 29, 42, 46, 47, 48, 49, 50, 51^, but CSF analytics is limited by the low volume obtained from single mice and by blood contaminations. The previous limitations are overcome by our subdural (“closed”) method for sampling murine CSF. The method allows collection of larger volumes (> 20 µl) of blood- and cell-free CSF, even upon repeated sampling and enables multiple protein analytical assays in parallel. We also provide an estimation of the absolute concentrations of hundreds of mouse CSF proteins by using the external UPS2 protein quantification standard.

The volume and purity of CSF gained from living mice are critical factors for accurate protein analytics and thus, for robust and reproducible insights into physiological and pathophysiological processes. A volume of 10-20 µl is often required for ELISA assays and is important when multiple protein analytical assays are to be run from single samples. A high purity of CSF is essential because contaminations with blood, which has a much higher protein concentration than CSF^22^, may profoundly distort CSF protein concentrations. Previously established CSF sampling techniques achieved volumes of 10-20 µl ^13, 15^, while our improved method allows routine sampling of 20 – 35 µl CSF from living mice. Larger volumes of up to 40 µl were only reported in cases, where CSF was collected post-mortem^12^ and, consequently, no longer reflected the CSF composition of living mice, or when the CSF contained extra-cerebrospinal impurities due to the impure nature of the CSF-collection technique by dura puncture and free flow of CSF over the dura (extradural or “open” collection) ^11, 52^.

Our improved method is based on the continuous CSF production in ventricles and its circulation through the CNS^8, 53,15^. We used a stepwise (every 4-6 minutes) low-grade negative pressure, applied only when the CSF pulsated again in the capillary (i.e. pressure equalization). This subdural (“closed”) collection approach avoided forced CSF suction and possible tissue damage, thus, resulting in a high CSF quality and enabled to collect more than 20 μl CSF within 35-45 minutes in comparison to a passive flow into the capillary, where a volume of approximately 8 μl of CSF fluid was obtained ^13, 14^.

Our study provides a rough estimation of the concentrations of hundreds of individual CSF proteins (in fmol/μl) based on an external calibration using a commercially available protein reference set covering a wide dynamic range and using intensity-based absolute quantification (iBAQ)^54, 55, 56^. This concentration estimation is simple, cost-efficient and easily feasible for all detected proteins in the murine CSF. The variations in protein concentration between murine and human CSF stem from species differences, as well as the method of cisternal CSF collection in mice versus lumbar CSF collection in humans. A precise concentration determination of individual proteins would require the much more expensive, absolutely quantified isotopically labeled spike-in peptides^57^, if using mass spectrometry, or other methods, such as immunoassays for which standard curves for each protein need to be determined. As such, the concentration estimation is particularly helpful when setting up subsequent targeted assays, such as immunoassays that have a limited detection range, or when comparing protein concentrations between preclinical animal species, between different experiments, between different laboratories or with humans.

Another outcome of our study is the result from the repeated CSF sampling in mice. We found that the minimal dural trauma due to the puncture for CSF collection induced reactive changes of proteins in the CSF that persisted up to 3 months after the first puncture despite macroscopical restoration of the cisterna magna and dura. The exact reason for the induced proteome changes is not known. Clinically, this may correlate to a widespread meningeal reaction after a single lumbar puncture in some patients, as indicated by reactive gadolinium enhancement of the cranial meninges even weeks after puncture^58^. The changes in CSF upon repeated sampling need to be taken into consideration for longitudinal studies in mice.

Finally, we investigated the CSF proteome of the widely used Alzheimer’s 5xFAD mouse model. At seven months of age, 5xFAD have a severe plaque pathology and microgliosis ^40^. Consequently, compared to wild-type mice, we identified significantly increased abundances of microglia-enriched proteins such as the cathepsins, Cd14, Csf1r, Ly86, Lag3 and the microglia activation marker Trem2. These microglial protein signatures are consistent with CSF proteome changes previously identified in another AD mouse model, the APPPS1 model^42^, but at a later time point. These findings are well in line with the much faster-progressing pathology in 5xFAD compared to APPPS1 mice. The largest fold-change was found for Neurofilament M (Nefm), a marker for neurodegeneration^3^. The other two neurofilaments, L and H, were only detected and quantified in the CSF of 5xFAD mice, but below the limit of quantification for the wild-type littermates. Similar changes of Nefm have also been observed in the CSF of 18-month-old APPPS1 and A30PαS mice^42^. Increased neurofilament abundance is seen as a sensitive marker for neurodegeneration, even before the massive loss of neurons that characterizes neurodegenerative disease. Thus, it is likely that the increased neurofilament levels in the CSF of 5xFAD mice at seven months indicates that neurodegeneration already starts at seven months or earlier, whereas overt neuronal loss has been previously detected at nine months, but was still absent at six months of age^41, 43^. Increased CSF levels of calbindin 2 (Calb2), also known as calretinin, might also be related to the degeneration of calretinin-positive interneurons, which has been described for 12-month-old 5xFAD mice^59^. According to single-cell RNA sequencing, calretinin is mainly expressed in inhibitory neurons^60^. Hence, CSF calretinin is a novel candidate as a neurodegeneration marker.

In conclusion, our optimized protocol facilitates the collection of large CSF volumes from mice with high quality, which offers the possibility to perform various biochemical analyses using individual CSF samples. This method will help to reduce the number of mice required for CSF studies and will also be instrumental in gaining important new insights into basic and applied neuroscience, such as into brain disorders, the consequences of trauma or drug treatments as well as the function of specific genes/proteins, such as genetic risk factors for brain diseases.

## Acknowledgements

We thank Anna Berghofer and Brigitte Nuscher for excellent technical support and Burcu Seker for providing the 5xFAD mice for this study. The study was supported by a Marie-Curie IntraEuropean Fellowship grant (FP7-PEOPLE-2013-IEF, project number 625970), by the Deutsche Forschungsgemeinschaft (DFG, German Research Foundation) under Germany’s Excellence Strategy within the framework of the Munich Cluster for Systems Neurology (EXC 2145 SyNergy– ID 390857198), by the BMBF through project CLINSPECT-M and by the Cure Alzheimer’s Fund.

## Author Contribution statement

AL, SAM, SFL and NP designed the study. AL and SAM developed the methods, performed experiments – with the help of GJ - and analyzed the data. AL and SL wrote the manuscript. All authors critically reviewed the manuscript; SL and NP supervised the present work.

## Materials and Methods

### Animal handling

Adult male C57BL/6 mice (3, 6, and 12 months old, Charles Rivers Laboratories, n = 36 in total) 5xFAD (7 months old, n=5), and its corresponding wild-type (7 months old, n=5) male littermate mice were used for the study. The 5xFAD mice model Alzheimer’s Disease amyloid pathology; they overexpress mutant human APP695 with the Swedish (K670N, M671L), Florida (I716V), and London (V717I) Familial Alzheimer’s Disease (FAD) mutations along with human PS1 harboring two FAD mutations, M146L and L286V^40^.

All animals were housed in our animal facility, under a 12/12hrs light/dark cycle and were provided food and water ad libitum. Experiments were conducted according to institutional guidelines of the University of Munich after approval of the Ethical Review Board of the Government of Upper Bavaria.

### Optimized CSF collection from cisterna magna - surgical protocol

Collection of the CSF was modified from previously published protocols^13^ ^15^. Initial setup experiments for procedure-standardization were performed on 3-month-old C57BL/6 mice (n = 6), which were not used for further analyses. We used the “subdural” method of CSF collection, with no contact to extradural tissues. We describe the protocol in detail below (Figure 1). The operative set-up and tools needed for the procedure are shown in Figure 1A and supplementary table 1 (with purchase codes). Mice are anesthetized with ketamine/xylazine (100/10mg per kg of body weight) and placed on a stereotactic frame with the head bent at approximately 135° (Figure 1B). The surgical procedure is shown in sequential steps (photos) in Figure 1C and 1D. An oil-based cream (such as the Bepanthol Eye-cream) is applied on the skin to pull the cervical fur to both sides of the incision line (Figure 1C, first photo). The skin at the incision line is locally sterilized using 70% alcohol with the aid of cotton-bud. The skin of dorsal upper cervical region is incised longitudinally for approximately 1cm starting from the base of the skull towards the 2^nd^ cervical vertebra, i.e. starting from a line between the ears downwards for 1cm (red dotted line in Figure 1C, step 1), thus revealing the occipital bone, the splenius capitis and the respective fascia (Figure 1C, step 2). Under a dissecting microscope, the superficial cervical muscles (m. splenius) and the thin covering fascia (black lines in Figure 1C, step 3) are carefully separated by forceps along and up to the median fibrous raphe of the neck (dotted black line in Figure 1C, step 3 and 4)^61^ without causing tissue bleeding. A sequential midline cut of approximately 0.5cm is done along the median fibrous raphe of the neck to further separate the splenius capitis and hence increase the surgical filed view (Figure 1C, step 5), taking always care to avoid vessel or muscle cutting which would cause local hemorrhage; here the semispinalis capitis is revealed, under of which the dura of cisterna magna will be found later (Figure 1C, dotted black triangle in step6). When necessary, hemostasis is performed by blunt pressure on the muscles, always before exposing the underlying cervical dura. Next, with the aid of the angled forceps and the self-made hooks (see Figure 1A) the muscles are pulled to the sides to open the surgical field, revealing the dura of the cisterna magna (see steps 7-11 following in Figure 1C). Initially the angled forceps is inserted just under the semispinalis muscles to guide the insertion of the left hook (Figure 1C step 7, position pointed by a black hooked arrow). By correct positioning of the forceps, the left hook is carefully inserted under the semispinalis capitis (Figure 1C step 8) and its sideward forced retraction reveals the left half part of the cisterna magna (Figure 1C step 9). Similarly, the angled forceps guides the insertion of the right hook (Figure 1C steps 9-11). Eventually, by pulling and attaching the strings of the hooks securely to the sides of the earpieces, the cisterna magna is non-traumatically revealed (Figure 1C step 12), ready for capillary puncture and CSF collection.

The exposed dura is punctured by a pulled glass capillary. The tip of the capillary has been already cut beveled by the forceps in an angle of approximately 45° (Figure 1D) under the optical control of the dissecting microscope, just before the glass capillary is connected to the tubing system for suction; the tubing system is a polyethylene PE90 tube connecting the 20 gauge needle (see Figure 1A) of the 1ml suction syringe to the glass capillary. The latter is carefully stabilized on the head of the stereotactic arm (Figure 1A) just before dura puncturing, to avoid accidental breaking of its fine tip. Once the dura is prepared, the capillary tip is then carefully advanced in position to puncture the dura at an angle of approximately 45° (Figure 1D, second photo): in order to avoid hemorrhages during puncturing, the tip has to point at the depth of the cisterna magna away of the dorsal spinal artery or its branches, as shown in Figure 1D (third photo). Once in position, the tip is further advanced to puncture the dura using the micromanipulator controls of the stereotaxic frame. Puncture of the dura by the capillary requires extra force to overcome the resistance of the dura; extra care is necessary to avoid tissue injury by abrupt or excessive tip advancement. Here, some additional head-bending before capillary positioning can stretch the dura and thus ease its puncture by the tip.

Once the tip is successfully inside the cisterna magna cavity (Figure 1D), CSF flows initially freely in the capillary. At that point, the capillary is slightly pulled back for approximately 0.4-0.6mm - under visual microscopic control- to gently pull the dura up and increase the volume of the cisterna magna, facilitating further flow of the CSF into the capillary. Still, if the capillary is pulled too much from the dura and cisterna magna the CSF will leak extradural and the procedure is considered failed due to contamination of the CSF.

An initial free-flow of CSF into the glass capillary reflects the normal opening CSF pressure and fills a column of approximately 12-15 mm in the capillary. In order to increase the collected CSF volume, the application of a mild stepwise vacuum via mild suction by the 1 ml syringe (at discrete steps of 20-30 μl every 4-5minutes) is necessary. Each suction step has to be applied a) when the CSF column does not rise anymore in the glass capillary and it pulsates within it, indicating a pressure equilibrium, and b) when the cisterna has partially refilled. Then, the suction step slightly overcomes the hydrostatic CSF pressure and allows for additional CSF withdrawal (verified by visible CSF withdrawal in the capillary). A partial collapse of the cisterna magna with caudal retraction of the cerebellum is observed and expected, here the black arrow in Figure 1D (fourth photo) points at the retraction of the cerebellum from the initial position -black dotted line- to the new one -white dotted line). It is of extreme importance to perform each suction step patiently, so as to allow for pressure equilibrium and prevent complete caudal sanction of the cerebellum with resulting central herniation and brainstem tissue damage.

After the operation the animals can be allowed to wake up and be returned to their cage. No animal died during or after the CSF collection process at any timepoint. In the present study, animals were either sacrificed or kept alive after CSF sampling, depending on the experimental design. When kept alive, all animals displayed normal activity in their cages approximately 1.5-2 hours after anesthesia.

### Experimental design

All animals were randomized for CSF collection prior to the surgery. Group codes were broken only after data validation and sample analysis. Group size for further MS proteomic analysis was calculated based on previous data. We initially setup and improved the surgical procedure of blood-free CSF collection (see above) in C57BL/6 mice (n=6), then we applied the method on wild type C57BL/6 and transgenic animals as described below. All experiments followed the same CSF collection and aliquoting procedure.

We used three experimental setups to show-case the validity of our concept: a) single CSF collections from separate 3, 6 and 12 old C57BL/6 mice (n= 9, 5, and 5 respectively), to study aging-related CSF changes, CSF purity and quantification possibilities, b) a first CSF collection at three months and a second (repeated) CSF collection from the same mice (n= 6), to show the feasibility of multiple, repeated CSF collection from the same mice, and c) CSF collection from 7-month-old transgenic (5xFAD) mice and their littermates (n= 5 each), to showcase the feasibility of multiple molecular and spectrometric analyses in the collected CSF, also including post-collection contamination control by sensitive hemoglobin-ELISA. In addition, we used pooled CSF from normal C57BL7/6 mice (3 months old, n=5) for dynamic light scattering (DLS) analysis, as described below. All data are reported according to the ARRIVE criteria^62^.

### Human CSF collection

Anonymized leftover material of human CSF from 8 subjects was provided by Institute of Laboratory Medicine, University Hospital Ludwig Maximilian University Munich. The subjects did not have a history of head injury, brain ischemia, infarction, hemorrhage, infection, inflammation or degenerative neurological disease within 6 months before the collection of CSF. Samples were collected non-traumatically without blood contamination. Further handling and centrifugation of CSF was performed using routine clinical procedures. After centrifugation of CSF at 800 g it was stored at -80°C until further processing.

### Post-collection processing of CSF samples with quality control analysis

The aforementioned procedure yields mouse CSF without macroscopic blood/cell contamination. An initial centrifugation at 10,000 rpm (2000 x g) for 10 minutes at 4 ^°^C was routinely performed to discard any possible cells/debris according to most available CSF handling protocols. Although no pellet was evident macroscopically after this centrifugation, the supernatant was always carefully removed and an undisturbed, “pellet” of approximately 2-3 μl was left behind (putative cells/debris). The supernatant was then immediately aliquoted and stored at -80 ^°^C until further analyzed, while the pellet was reconstituted with PBS to a final volume of 10 μl, where necessary, for further microscopical analysis and quality control (QC).

The QC analysis of collected CSF was performed with the aid of a Neubauer chamber. The samples used were the CSF-pellets, reconstituted with PBS at 10μl of volume. As such, depending on the initially collected CSF volume (20 to 30 μl), this represented an approximate 2-3x concentration factor compared to the initial CSF volume. We measured the number of cells (N) in the total 25 small Neubauer squares (equal to 0.1μl of Neubauer volume) under 40x, using routine methodology. The number of cells per μl of collected CSF was then calculated using the formula: “cells/μl = (N x 10) / concentration factor”.

### Simulation experiments of blood contamination

In order to understand how a blood contamination of the CSF would change its microscopical image under the Neubauer chamber, we simulated this via diluted blood in PBS, at 1:100 and 1:10,000 dilutions, representing 1% and 0.01% contaminations respectively. Specifically, the latter dilution equals a cut-off limit previously accepted as “tolerable” in human CSF studies^22^. Here, we measured the number of erythrocytes (red-blood cells, RBC) using the routine RBC-methodology (in 5 small squares in Neubauer) and calculated the cells/ μl using the formula “cells/μl = N x 5 x 10”. All results (CSF and blood) are reported as “cells/ μl“. All samples were also analyzed for hemoglobin using an ELISA method (see below). Pure mouse plasma from 5 normal C57BL/6 adult mice was used as reference control for hemoglobin content.

### Dynamic Light Scattering analysis of particle size in collected CSF

Microparticle analysis of mouse and human CSF was performed with the aid of Dynamic light scattering (DLS)^63, 64, 65^. For the analysis we used a Zetasizer Nano S particle analyzer (Malvern Instruments Ltd., Malvern UK) and the corresponding software (Zetasizer 7.03). Solvent-resistant disposable micro cuvettes (UV-Cuvette micro 70μl, Brand GmbH, Germany, Cat. No. 759200) were used for experiments with a minimum sample volume of 40 µL. The measurements were made at a fixed position with an automatic attenuator and at a controlled temperature. While human CSF volume was adequate for DLS measurements, for mouse analyses we had to collect CSF from extra mice (C57BL/6, n=5) and use all volumes solely for this purpose. All CSF samples were measured before and after centrifugation at 800 and 2000 g and diluted at 1:5 with NaCl 0.9% to reach the required volume of 40 μl per cuvette. For each sample, 3 sets of 10 measurements were automated averaged to provide one average curve result for each sample. Peaks are expressed in nm +/- std. A calibration of the system was done with nanoparticles of 125 nm diameter (Suppl. Figure 2C).

### Processing of the mouse and human CSF samples for Liquid chromatography Mass Spectrometry analysis (LCMS)

Proteolytic digestion of 5 µl CSF aliquots was performed in 50 mM ammonium bicarbonate with 0.1% sodium deoxycholate. Disulfide bonds were reduced and cysteine residues were alkylated by addition of 2 µl 10 mM dithiothreitol (Biomol, Germany) and subsequently 2 µl 55 mM iodoacetamide (Sigma Aldrich, Germany). Proteolytic digestion was performed by consecutive digestion with LysC (0.1 µg; 4 h) and trypsin (0.1 µg; 16 h) at room temperature (Promega, Germany).

A volume of 4 µl of 8% formic acid (Sigma Aldrich, Germany) and 150 µl of 0.1% formic acid (Sigma Aldrich Germany) was added to acidify the samples. Precipitated deoxycholate was removed by centrifugation at 16,000 g for 10 min at 4 °C. Peptides were desalted by stop and go extraction (STAGE) with C18 tips. Samples were dried by vacuum and dissolved in 20 µL 0.1% formic acid in a sonication bath.

Eight out of 20 µL, correlating with a starting amount of 2 µL CSF, were separated on a nanoLC system (EASY-nLC 1000, Proxeon – part of Thermo Scientific, US) equipped with a PRSO-V1 column oven (Sonation, Germany) using an in-house packed C18 column (30 cm x 75 µm ID, ReproSil-Pur 120 C18-AQ, 1.9 µm, Dr. Maisch GmbH, Germany) with a binary gradient of water (A) and acetonitrile (B) containing 0.1% formic acid at 50°C column temperature and a flow of 250 nl/min (0 min, 2% B; 3:30 min, 5% B; 137:30 min, 25% B; 168:30 min, 35% B; 182:30 min, 60% B; 185 min, 95% B; 200 min, 95% B). The nanoLC was coupled online via a nanospray flex ion source (Proxeon – part of Thermo Scientific, US) to a Q-Exactive mass spectrometer (Thermo Scientific, US). Full MS spectra were acquired at a resolution of 70,000 (AGC target: 3×10^6^, mass range: 300-1400 m/z). The top 10 peptide ions exceeding an intensity of 5×10^4^ were chosen for collision induced dissociation (Resolution: 17,500, AGC target: 1×10^5^, isolation window: 2 m/z). A dynamic exclusion of 120 s was used for peptide fragmentation.

CSF samples from 5xFAD and wild-type control mice were analyzed on an EASY-nLC 1000 coupled to a Q-Exactive HF mass spectrometer (Thermo Scientific, US) equipped with a PRSO-V1 column oven (Sonation, Germany). Peptide separation was performed on the same columns using peptide separation applying a binary gradient of water and 80% acetonitrile (B) for 120 min at a flow rate of 250 nL/min and a column temperature of 50 °C: 3% B 0 min; 6% B 2 min; 30% B 92 min; 44% B 112 min; 75% B 121 min. Samples were analyzed using data dependent acquisition (DDA) and data independent acquisition (DIA) injecting 8 out of 20 µL. For DDA, full MS spectra were acquired at a resolution of 120,000 (AGC target: 3×10^6^, mass range: 300-1400 m/z). The top 15 peptide ions exceeding an intensity of 5×10^4^ were chosen for collision induced dissociation (Resolution: 15,000, AGC target: 1×10^5^, IT: 100ms, isolation window: 1.6 m/z). A dynamic exclusion of 120 s was used for peptide fragmentation. For DIA, one MS1 full scan was followed by 20 sequential DIA windows with variable width for peptide fragment ion spectra with an overlap of 1 m/z covering a scan range of 300 to 1400 m/z. Full scans were acquired with 120,000 resolution and an AGC target of 5x10^6^. Afterwards, 20 DIA windows were scanned with a resolution of 30,000 and an AGC of 3x10^6^. The maximum IT for fragment ion spectra was set to auto to achieve optimal cycle times.

### LCMS data analysis of CSF samples

Data from DDA runs were analyzed with Maxquant software (maxquant.org, Max-Planck Institute Munich)^66^ version 1.5.5.1 or version 2.1.4.0 for 5xFAD CSF. The MS data were searched against a reviewed canonical fasta database of Mus musculus from UniProt (download: March 9th 2017, 16851 entries) or a database with one protein sequence per gene from UniProt (download: January 12^th^ 2023, 16851 entries) for 5xFAD CSF. Trypsin was defined as protease. Two missed cleavages were allowed for the database search. The option first search was used to recalibrate the peptide masses within a window of 20 ppm. For the main search, peptide and peptide fragment mass tolerances were set to 4.5 and 20 ppm, respectively. Carbamidomethylation of cysteine was defined as static modification. Acetylation of the protein N-term as well as oxidation of methionine were set as variable modifications. The false discovery rate for both, peptides and proteins, was adjusted to less than 1% using a target and decoy approach (concatenated forward/reverse database). Only unique peptides were used for quantification. Label-free quantification (LFQ) of proteins required at least two ratio counts of unique peptides. Proteins were considered identified if they were detected by at least 2 unique peptides in at least 3 out of n sample of each group, based on LFQ values. The "intensity based absolute quantification" or iBAQ values were also calculated as previously described^56^. We use LFQ values for relative protein quantification among samples and iBAQ values for estimating absolute quantification values, as defined previously^55, 67^.

Data from DIA runs of the 5xFAD experiment were analyzed using the software DIA-NN version 1.8. A library free search was used with the same fasta database including a database of common contaminants such as trypsin and human keratins. Oxidation methionines and acetylation of protein N-termini were set as variable modifications, whereas carbamidomethylation of cyteines was set as fixed modification. Charge states from two to four with a m/z range of 300 to 1400 were considered. Mass accuracy settings were set to automatic. The match between runs option was applied. Data normalization was disabled.

We also take into consideration that LFQ values are normalized among samples while iBAQ values are not. Imputation of data was performed with the software perseus version 1.6.14.0 ^68^ in the cases of 5xFAD vs wild-type (wt) animal analyses to increase the visibility of the 5xFAD pathology versus the wt animals. Only protein groups with a complete quantification profile in at least one experimental group (5xFAD or wt) were considered and missing LFQ data was imputed after log2 transformation according to a normal distribution with a width of 0.3 and a down-shift of 1.8.

We refer to "qualitative changes" of proteins when we study the presence or absence of a protein in all samples of a group. To report "quantitative changes" we use LFQ values (for relative protein changes between 2 groups) or iBAQ values (for absolute quantification of proteins in samples) only when a protein is detected in both compared groups. Volcano plots are used to illustrate relative protein changes (LFQ values) between the experimental groups. A two-sided non-paired Student’s t-test was used to evaluate the significance of each protein change between two independent groups. For repeated sampling of the same animals at 3 and 6 months (3moR and 6moR groups), we used correspondingly a paired sample t-test. For absolute quantification of selected proteins in different samples (mouse or human) we use the log10 transformed iBAQ (intensity based absolute quantification) values, data that are below detection limit are presented in all graphs as "zero values".

### Estimation of CSF protein concentrations using the Proteomics Dynamic Range Standard UPS2 in LCMS

For the estimation of absolute CSF protein concentrations, we used the UPS2 kit proteomics dynamic range standard set, which covers 5 orders of magnitude, and performed a calibration based on the iBAQ intensities.

Briefly, the UPS2 standard was dissolved in 20 µL of 50 mM ammonium bicarbonate and 0.1% sodium deoxycholate. The UPS2 standard was digested with the same protocol as the CSF samples. Finally, an amount of 0.5% and 1% of the digested UPS2 standard, which covers a range from 500 to 0.0025 fmol (or 2.5 amol), was analyzed using the same LC-MS/MS method as above. The known amounts of different proteins were used to generate a calibration curve of absolute protein amounts in fmol on the basis of their iBAQ intensities, using a non-linear fit-analysis of data with the least-squares regression (see statistical analysis section). Data outliers were checked and mathematically excluded where necessary before generation of the calibration curves using the established ROUT method^69^ in GraphPad 9.

### Bioinformatics analysis

In order to identify proteins and relative biological functions that are enriched in the CSF samples we used the DAVID 6.8 Bioinformatics Resources of NIAID/NIH (https://david.ncifcrf.gov/summary.jsp)^70^. As indicated by the DAVID Database, we used the Functional Annotation Clustering to search for enrichment and consequently for major biological functions and pathways standing out in each of our samples in comparison to background mus musculus proteome. For this purpose, we checked for enrichment of different functional categories (UP-Keywords; Gene Ontology: GOTERM_CC/BP/MF_DIRECT; Pathways: KEGG and REACTOME)^71^ separately for proteins with significantly increased and decreased abundance.

Medium classification stringency, similarity term overlap of 3, similarity threshold of 0.5 for Kappa statistics and Enrichment Threshold (EASE) at 0.05 was applied. We selected and report the significant different protein clusters based on Benjamini p-values of p<0.05 enriched for "increased" and "decreased" proteins. The Benjamini p-values are adjusted for multiple comparisons to lower the family-wise false discovery rate and, thus, are more conservative than Fisher Exact p values.

### Enzyme-linked immunosorbent Assay of selected proteins (Hemoglobin, TREM2, NfL, ApoE and sAPP) in the mouse or human CSF

For detection of murine hemoglobin in mouse CSF (where applicable), mouse plasma and simulation experiments of blood contamination (see above), all samples were analyzed with a sensitive ELISA-kit (#ab157715, Abcam) according to the manufacturer’s instructions and as previously described^72^. Briefly, 1 µl of murine CSF was diluted 1:200 and measured in duplicates (of 0.5μl each); for plasma and simulation experiments we also used 1μl of sample. The computation of standard curves, as well as the analysis of samples with unknown concentrations, was conducted employing a four-parameter logistic fit curve, utilizing the resources available on myassays.com.

For detection of murine TREM2 in CSF of adult C57/BL6 wildtype or 5xFAD mice, the CSF was analyzed using a previously developed ELISA that was set up on the Meso Scale Discovery (MSD) platform as described previously^44^. Briefly, a streptavidin-coated small-spot 96-well plate (MSD, #L45SA-2) was incubated overnight in blocking solution (PBS, 0.05% Tween, 3% BSA) at 4°C, before the plate was incubated with 25 µl of a biotinylated-anti-TREM2 antibody (#BAF1729, biotechne) at a concentration of 0.125 µg/ml for 90 min. The plate was rotated horizontally on a shaker at 300 rpm. After this, the following incubation steps were carried out at room temperature using a horizontal shaker at 300 rpm. The plate was washed 3x with 250 µl of washing buffer (PBS, 0.05% Tween) per well. Standards were prepared in a two-fold serial dilution using recombinant TREM2-FC (#1729-T2, biotechne) ranging from 400 – 12.5 pg/ml. Prior to incubation, the TREM2 standards were denatured by addition of a denaturation buffer (final conc.: 20 mM Tris-HCL pH 6.8, 0.4% SDS, 4% Glycerol, 0.2% ß-ME, 5 mM EDTA) and boiling for 5 min at 95°C. The collected CSF was diluted 1:5 in dilution buffer (PBS, 0.05% Tween, 1% BSA, 2 µl/ml protease inhibitor cocktail freshly added). 50 µl of standards and samples were dispensed in the wells and incubated for 120 min. The plate was cleansed mentioned above, followed by the addition of 50 µl of a 1 µg/ml rat anti-TREM2 detection antibody (#MABN2321, Sigma Aldrich) into the wells, which was left to incubate for 60 min. The plate underwent another round of the previously detailed washing process, after which a goat-anti-rat Sulfo-tag (#R32AH-1, MSD) secondary antibody was introduced into the wells. This secondary antibody was diluted 1:1000 in blocking buffer, with 25 µl of the mixture added to each well, followed by a 60 min incubation period. Following this, the plate was washed twice with washing buffer and twice with PBS, and then 150 µl of a 1x Read buffer (MSD) was poured into the wells. The plate’s contents were then read utilizing the internal MSD platform. To calculate the TREM2 levels in the unknown concentration samples, the MSD platform software was employed, utilizing a four-parameter logistic fit curve regression model.

The concentration of NfL in our 5xFAD mouse CSF samples was measured using the NF-light Advantage Assay Kit (Item 103186) for the Single Molecule Array (SIMOA) immunoassay SR-X (Quanterix). The analysis was carried out following the manufacturer’s protocol. Briefly, 3.4 µl CSF of adult C57/BL6 wildtype or 5xFAD mice was used to determine the levels of Nfl. The computation of standard curves, as well as the analysis of samples with unknown concentrations, was conducted employing a four-parameter logistic fit curve, utilizing the default settings of the software provided by Quanterix.

ApoE and sAPP were measured in human CSF samples using the Apolipoprotein E Human ELISA Kit (#EHAPOE) and Amyloid Precursor Protein Human ELISA Kit (#KHB0051) kits respectively from Life Technologies, following the manufactureŕs protocol in duplicates using a 1:100 dilution of CSF.

### Statistical analysis

Statistical analysis and graphs are performed with the GraphPad Prism 9. Venn diagrams were constructed using the online tool InteractiVenn^73^. Analyses of selected iBAQ values between groups were performed with appropriate parametric or nonparametric tests and post-hoc tests adjusted for multiple comparisons. Repeated data for 3moR and 6moR groups were analyzed with paired samples t-test. The UPS2 fmol-iBAQ curve analysis and construction was performed using the Non-linear Fit analysis of data (for a log-log X-Y line) with the least squares regression with appropriate weighting method, to best fit the curve data. Data in text are reported as mean±SD, unless noted differently; plotted data in graphs are shown as means ± 95% Confidence Interval (± 95%CI, unless noted differently) with superimposition of single values where necessary. Level of significance is set at 0.05 for all statistics.

## References

1. Teunissen CE, et al. White paper by the Society for CSF Analysis and Clinical Neurochemistry: Overcoming barriers in biomarker development and clinical translation. Alzheimers Res Ther 10, 30 (2018).

2. Olsson B, et al. CSF and blood biomarkers for the diagnosis of Alzheimer’s disease: a systematic review and meta-analysis. Lancet Neurol 15, 673–684 (2016).

3. Yuan A, Nixon RA. Neurofilament Proteins as Biomarkers to Monitor Neurological Diseases and the Efficacy of Therapies. Front Neurosci 15, 689938 (2021).

4. Simon MJ, Iliff JJ. Regulation of cerebrospinal fluid (CSF) flow in neurodegenerative, neurovascular and neuroinflammatory disease. Biochim Biophys Acta 1862, 442–451 (2016).

5. Wright BL, Lai JT, Sinclair AJ. Cerebrospinal fluid and lumbar puncture: a practical review. J Neurol 259, 1530–1545 (2012).

6. Dislich B, et al. Label-free Quantitative Proteomics of Mouse Cerebrospinal Fluid Detects beta-Site APP Cleaving Enzyme (BACE1) Protease Substrates In Vivo. Mol Cell Proteomics 14, 2550–2563 (2015).

7. Smith JS, Angel TE, Chavkin C, Orton DJ, Moore RJ, Smith RD. Characterization of individual mouse cerebrospinal fluid proteomes. Proteomics 14, 1102–1106 (2014).

8. Johanson CE, Duncan JA, 3rd, Klinge PM, Brinker T, Stopa EG, Silverberg GD. Multiplicity of cerebrospinal fluid functions: New challenges in health and disease. Cerebrospinal Fluid Res 5, 10 (2008).

9. Rudick RA, Zirretta DK, Herndon RM. Clearance of albumin from mouse subarachnoid space: a measure of CSF bulk flow. J Neurosci Methods 6, 253–259 (1982).

10. Bacioglu M, et al. Neurofilament Light Chain in Blood and CSF as Marker of Disease Progression in Mouse Models and in Neurodegenerative Diseases. Neuron 91, 56–66 (2016).

11. Maia LF, et al. Changes in amyloid-beta and Tau in the cerebrospinal fluid of transgenic mice overexpressing amyloid precursor protein. Sci Transl Med 5, 194re192 (2013).

12. Sakic B. Cerebrospinal fluid collection in laboratory mice: Literature review and modified cisternal puncture method. J Neurosci Methods 311, 402–407 (2019).

13. Liu L, Duff K. A technique for serial collection of cerebrospinal fluid from the cisterna magna in mouse. J Vis Exp, (2008).

14. Liu L, Herukka SK, Minkeviciene R, van Groen T, Tanila H. Longitudinal observation on CSF Abeta42 levels in young to middle-aged amyloid precursor protein/presenilin-1 doubly transgenic mice. Neurobiol Dis 17, 516–523 (2004).

15. Lim NK, Moestrup V, Zhang X, Wang WA, Moller A, Huang FD. An Improved Method for Collection of Cerebrospinal Fluid from Anesthetized Mice. J Vis Exp, (2018).

16. Cunningham R, Jany P, Messing A, Li L. Protein changes in immunodepleted cerebrospinal fluid from a transgenic mouse model of Alexander disease detected using mass spectrometry. J Proteome Res 12, 719–728 (2013).

17. Liddelow SA, et al. Cellular specificity of the blood-CSF barrier for albumin transfer across the choroid plexus epithelium. PLoS One 9, e106592 (2014).

18. Whish S, et al. The inner CSF-brain barrier: developmentally controlled access to the brain via intercellular junctions. Front Neurosci 9, 16 (2015).

19. Deisenhammer F, et al. Guidelines on routine cerebrospinal fluid analysis. Report from an EFNS task force. Eur J Neurol 13, 913–922 (2006).

20. Fogh JR, Jacobsen AM, Nguyen T, Rand KD, Olsen LR. Investigating surrogate cerebrospinal fluid matrix compositions for use in quantitative LC-MS analysis of therapeutic antibodies in the cerebrospinal fluid. Anal Bioanal Chem 412, 1653–1661 (2020).

21. Aasebo E, Opsahl JA, Bjorlykke Y, Myhr KM, Kroksveen AC, Berven FS. Effects of blood contamination and the rostro-caudal gradient on the human cerebrospinal fluid proteome. PLoS One 9, e90429 (2014).

22. Barkovits K, et al. Blood Contamination in CSF and Its Impact on Quantitative Analysis of Alpha-Synuclein. Cells 9, (2020).

23. Pigoni M, et al. Seizure protein 6 and its homolog seizure 6-like protein are physiological substrates of BACE1 in neurons. Molecular neurodegeneration 11, 67 (2016).

24. Tushaus J, et al. An optimized quantitative proteomics method establishes the cell type-resolved mouse brain secretome. EMBO J 39, e105693 (2020).

25. Emelyanov A, et al. Cryo-electron microscopy of extracellular vesicles from cerebrospinal fluid. PLoS One 15, e0227949 (2020).

26. Wahrle SE, et al. ABCA1 is required for normal central nervous system ApoE levels and for lipidation of astrocyte-secreted apoE. J Biol Chem 279, 40987–40993 (2004).

27. Tsujita M, et al. Apolipoprotein A-I in mouse cerebrospinal fluid derives from the liver and intestine via plasma high-density lipoproteins assembled by ABCA1 and LCAT. FEBS Lett 595, 773–788 (2021).

28. Ribeiro CA, et al. Transthyretin stabilization by iododiflunisal promotes amyloid-beta peptide clearance, decreases its deposition, and ameliorates cognitive deficits in an Alzheimer’s disease mouse model. J Alzheimers Dis 39, 357–370 (2014).

29. Jiang R, et al. Increased CSF-decorin predicts brain pathological changes driven by Alzheimer’s Abeta amyloidosis. Acta Neuropathol Commun 10, 96 (2022).

30. Nordestgaard LT, Christoffersen M, Frikke-Schmidt R. Shared Risk Factors between Dementia and Atherosclerotic Cardiovascular Disease. Int J Mol Sci 23, (2022).

31. Fornasiero EF, et al. Precisely measured protein lifetimes in the mouse brain reveal differences across tissues and subcellular fractions. Nat Commun 9, 4230 (2018).

32. Hubner NC, Nguyen LN, Hornig NC, Stunnenberg HG. A quantitative proteomics tool to identify DNA-protein interactions in primary cells or blood. J Proteome Res 14, 1315–1329 (2015).

33. Soufi B, Krug K, Harst A, Macek B. Characterization of the E. coli proteome and its modifications during growth and ethanol stress. Front Microbiol 6, 103 (2015).

34. Chang C, et al. PANDA: A comprehensive and flexible tool for quantitative proteomics data analysis. Bioinformatics 35, 898–900 (2019).

35. Zhang X, et al. An Interaction Landscape of Ubiquitin Signaling. Mol Cell 65, 941–955 e948 (2017).

36. Geiger T, Cox J, Mann M. Proteomics on an Orbitrap benchtop mass spectrometer using all-ion fragmentation. Mol Cell Proteomics 9, 2252–2261 (2010).

37. Koch S, et al. Characterization of four lipoprotein classes in human cerebrospinal fluid. J Lipid Res 42, 1143–1151 (2001).

38. Yamauchi K, et al. Apolipoprotein E in cerebrospinal fluid: relation to phenotype and plasma apolipoprotein E concentrations. Clin Chem 45, 497–504 (1999).

39. Cuchillo-Ibanez I, et al. Heteromers of amyloid precursor protein in cerebrospinal fluid. Mol Neurodegener 10, 2 (2015).

40. Oakley H, et al. Intraneuronal beta-amyloid aggregates, neurodegeneration, and neuron loss in transgenic mice with five familial Alzheimer’s disease mutations: potential factors in amyloid plaque formation. J Neurosci 26, 10129–10140 (2006).

41. Zhang L, Li J, Lin A. Assessment of neurodegeneration and neuronal loss in aged 5XFAD mice. STAR Protoc 2, 100915 (2021).

42. Eninger T, et al. Signatures of glial activity can be detected in the CSF proteome. Proc Natl Acad Sci U S A 119, e2119804119 (2022).

43. Eimer WA, Vassar R. Neuron loss in the 5XFAD mouse model of Alzheimer’s disease correlates with intraneuronal Abeta42 accumulation and Caspase-3 activation. Mol Neurodegener 8, 2 (2013).

44. Kleinberger G, et al. TREM2 mutations implicated in neurodegeneration impair cell surface transport and phagocytosis. Sci Transl Med 6, 243ra286 (2014).

45. Richter F, Meurers BH, Zhu C, Medvedeva VP, Chesselet MF. Neurons express hemoglobin alpha- and beta-chains in rat and human brains. J Comp Neurol 515, 538–547 (2009).

46. Tushaus J, et al. The pseudoprotease iRhom1 controls ectodomain shedding of membrane proteins in the nervous system. FASEB J 35, e21962 (2021).

47. Sleat DE, et al. Analysis of Brain and Cerebrospinal Fluid from Mouse Models of the Three Major Forms of Neuronal Ceroid Lipofuscinosis Reveals Changes in the Lysosomal Proteome. Mol Cell Proteomics 18, 2244–2261 (2019).

48. Wong JK, et al. Apolipoprotein B-100-mediated motor neuron degeneration in sporadic amyotrophic lateral sclerosis. Brain Commun 4, fcac207 (2022).

49. Greco F, et al. Longitudinal Bottom-Up Proteomics of Serum, Serum Extracellular Vesicles, and Cerebrospinal Fluid Reveals Candidate Biomarkers for Early Detection of Glioblastoma in a Murine Model. Molecules 26, (2021).

50. Herzog DP, et al. Longitudinal CSF proteome profiling in mice to uncover the acute and sustained mechanisms of action of rapid acting antidepressant (2R,6R)-hydroxynorketamine (HNK). Neurobiol Stress 15, 100404 (2021).

51. Zlatic SA, et al. Convergent cerebrospinal fluid proteomes and metabolic ontologies in humans and animal models of Rett syndrome. iScience 25, 104966 (2022).

52. DeMattos RB, et al. Plaque-associated disruption of CSF and plasma amyloid-beta (Abeta) equilibrium in a mouse model of Alzheimer’s disease. J Neurochem 81, 229–236 (2002).

53. Iliff JJ, Nedergaard M. Is there a cerebral lymphatic system? Stroke 44, S93–95 (2013).

54. Krey JF, et al. Accurate label-free protein quantitation with high- and low-resolution mass spectrometers. J Proteome Res 13, 1034–1044 (2014).

55. Ankney JA, Muneer A, Chen X. Relative and Absolute Quantitation in Mass Spectrometry-Based Proteomics. Annu Rev Anal Chem (Palo Alto Calif) 11, 49–77 (2018).

56. Schwanhausser B, et al. Global quantification of mammalian gene expression control. Nature 473, 337–342 (2011).

57. Lai X, Wang L, Witzmann FA. Issues and applications in label-free quantitative mass spectrometry. Int J Proteomics 2013, 756039 (2013).

58. Hannerz J, Ericson K, Bro Skejo HP. MR imaging with gadolinium in patients with and without post-lumbar puncture headache. Acta Radiol 40, 135–141 (1999).

59. Giesers NK, Wirths O. Loss of Hippocampal Calretinin and Parvalbumin Interneurons in the 5XFAD Mouse Model of Alzheimer’s Disease. ASN Neuro 12, 1759091420925356 (2020).

60. Karlsson M, et al. A single-cell type transcriptomics map of human tissues. Sci Adv 7, (2021).

61. Ruberte Ja, Carretero Aa, Navarro Ma. Morphological mouse phenotyping : anatomy, histology and imaging. Editorial Medica Panamericana S.A. (2017).

62. Kilkenny C, Browne WJ, Cuthill IC, Emerson M, Altman DG. Improving bioscience research reporting: the ARRIVE guidelines for reporting animal research. PLoS Biol 8, e1000412 (2010).

63. Stetefeld J, McKenna SA, Patel TR. Dynamic light scattering: a practical guide and applications in biomedical sciences. Biophys Rev 8, 409–427 (2016).

64. Palmieri V, et al. Dynamic light scattering for the characterization and counting of extracellular vesicles: a powerful noninvasive tool. J Nanopart Res 16, (2014).

65. Hallett FR, Watton J, Krygsman P. Vesicle sizing: Number distributions by dynamic light scattering. Biophys J 59, 357–362 (1991).

66. Cox J, Hein MY, Luber CA, Paron I, Nagaraj N, Mann M. Accurate proteome-wide label-free quantification by delayed normalization and maximal peptide ratio extraction, termed MaxLFQ. Mol Cell Proteomics 13, 2513–2526 (2014).

67. Megger DA, Bracht T, Meyer HE, Sitek B. Label-free quantification in clinical proteomics. Biochim Biophys Acta 1834, 1581–1590 (2013).

68. Tyanova S, Cox J. Perseus: A Bioinformatics Platform for Integrative Analysis of Proteomics Data in Cancer Research. *Methods in molecular biology (Clifton*, NJ) 1711, 133–148 (2018).

69. Motulsky HJ, Brown RE. Detecting outliers when fitting data with nonlinear regression - a new method based on robust nonlinear regression and the false discovery rate. BMC Bioinformatics 7, 123 (2006).

70. Huang da W, Sherman BT, Lempicki RA. Systematic and integrative analysis of large gene lists using DAVID bioinformatics resources. Nat Protoc 4, 44–57 (2009).

71. Hauck SM, et al. Deciphering membrane-associated molecular processes in target tissue of autoimmune uveitis by label-free quantitative mass spectrometry. Mol Cell Proteomics 9, 2292–2305 (2010).

72. Simoes S, et al. Tau and other proteins found in Alzheimer’s disease spinal fluid are linked to retromer-mediated endosomal traffic in mice and humans. Sci Transl Med 12, (2020).

73. Heberle H, Meirelles GV, da Silva FR, Telles GP, Minghim R. InteractiVenn: a web-based tool for the analysis of sets through Venn diagrams. BMC Bioinformatics 16, 169 (2015).

