## Supplement for "An improved method for sampling and quantitative protein analytics of cerebrospinal fluid of single mice"

**Supplementary Table 1: Suggested equipment for CSF collection from cisterna magna.** Ordering numbers in brackets indicate suggested but not mandatory ordering codes or companies.

| Equipment | Description | Supplier | Ordering No. |
|---|---|---|---|
| **Needle 20 Gauge** | needle 20 Gauge for PE tube connection | any available | NA |
| **Tubing, PE-90/10 (1.27OD x 0.86ID)** | Polyethylene Tubing (3m) | Warner Instruments | 64-0754 |
| **1ml syringe** | 1ml syringe | any available | NA |
| **insulin needles (Gauge 25)** | insulin needles (Gauge 25) for self-made hooks (recommended 4 hooks) | any available | NA |
| **Dumont #5/45 Forceps** | FST Dumont Forceps #5/45 Angled 45o tip 0.1 x 0.06mm | Fine Science Tools | 11251-35 |
| **Dumont Forceps Dumoxel** | FST Dumont #3 Forceps | Fine Science Tools | 11231-30 |
| **Bonn Scissors** | Straight sharp/sharp tip, 9cm length scissor | Fine Science Tools | 14184-09 |
| **Stereotaxic Frame for Mice** | Just for Mouse Stereotaxic Instrument (or other equivalent) | Stoelting (or other equivalent) | (51730) |
| **Heating pad** | Heating pad for stereotaxic frame | Stoelting (or other equivalent) | (53850M) |
| **Eppendorf Protein LoBind tubes 0.5ml** | 0.5 mL tubes with low protein binding | Sigma-Aldrich (or other equivalent) | (EP0030108094) |
| **Glass capillaries** | Borosilicate Glass OD 1.0mm, ID 0.75mm, 10cm length | Sutter Instrument | B100-75-10 |
| **Capillary puller** | P-1000 Micropipette Puller (or other equivalent) | (Sutter Instrument) | (P-1000) |
| **Ketamin-xylazin** | ketamin-xylazin 100:10 mg/kg mixture | any available | NA |

**Supplementary Table 2: Estimated concentrations of abundant proteins in normal human CSF** (n=8), as well as proteins at iBAQ range of $10^7$. NA = data not available; "values in literature" indicate (where present) the reported range of mean± SD (or SE according to the reference), for Hemoglobin data "<...>" indicates values that have to be interpreted with caution.

| Protein ID | Protein names (human) | Gene names | iBAQ value (± SD) | Estimated values in fmol/µL (± SD) | Estimated values in mg/l (± SD) | Values in literature (mean ± SD or SE) |
|---|---|---|---|---|---|---|
| **P02766** | Transthyretin | TTR | 1.57E+09 ± 1.47E+08 | 193.1 ± 24.5 | 3.1±0.4 mg/l | 15.8 (5.8-26.1) mg/l [73] |
| **P02649** | Apolipoprotein E | APOE | 1.48E+09 ± 4.56E+08 | 181.4 ± 61.4 | 6.5±2.2 mg/l | 3-9 ± 2-3 mg/l [37, 38] |
| **P02787** | Serotransferrin | TF | 7.23E+08 ± 1.64E+08 | 79.8 ± 20.6 | 6.1±1.6 mg/l | NA |
| **P10909** | Clusterin | CLU | 8.45E+08 ± 2.58E+08 | 95.8 ± 32.4 | 5.0±1.7 mg/l | 1.9 ± 0.036 mg/l [74] |
| **P01024** | Complement 3 | C3 | 2.33E+08 ± 6.37E+07 | 21.9 ± 6.8 | 4.1±1.3 mg/l | 1.8 (0.8-3.3) mg/l [75] |
| **P00441** | Superoxide dismutase [Cu-Zn] | SOD1 | 7.14E+07 ±3.26E+07 | 5.73 ±2.99 | 0.09±0.05 mg/l | 0.135 ± 0.047 mg/l [76] |
| **P05067** | Amyloid beta A4 protein | APP | 1.00E+08 ±3.64E+07 | 8.42 ± 3.42 | 0.73±0.3 mg/l | 0.75 mg/l [39] |
| **P31946** | 14-3-3 protein β/α | YWHAB | 2.41E+05 ± 9.06E+04 | 0.009 ±0.004 | 0.2±0.1 µg/l | NA for healthy subjects |
| **P01031** | Complement C5 | C5 | 8.23E+06 ± 5.31E+06 | 0.49 ± 0.36 | 0.09±0.07 mg/l | Approx. 0.2±0.1 mg/l [77] |
| **P02790** | Hemopexin | HPX | 9.10E+08± 2.00E+08 | 103.8±25.9 | 5.4±1.3 mg/l | 22.6 mg/l [78] |
| **P13521** | Secretogranin-2 | SCG2 | 2.23E+07 ± 9.62E+06 | 1.51±0.72 | 0.11±0.05 mg/l | Approx. 1.8 fmol/µl [79] |
| **Q02818** | Nucleobindin-1 | NUCB1 | 9.07E+06 ± 2.38E+06 | 0.54±0.16 | 0.03±0.01 mg/l | NA |
| **Q99983** | Osteomodulin | OMD | 1.23E+07 ± 6.52E+06 | 0.77±0.47 | 0.04±0.02 mg/l | NA |
| **Q06828** | Fibromodulin | FMOD | 5.94E+06 ± 2.33E+06 | 0.33±0.15 | 0.014±0.006 mg/l | NA |
| **P68871** | Hemoglobin subunit beta | HBB; HBD | 1.00E+07 ±2.27E+07 | 0.74 ±1.77 | 0.012±0.028 mg/l | *<3mg/dl>* [80] |

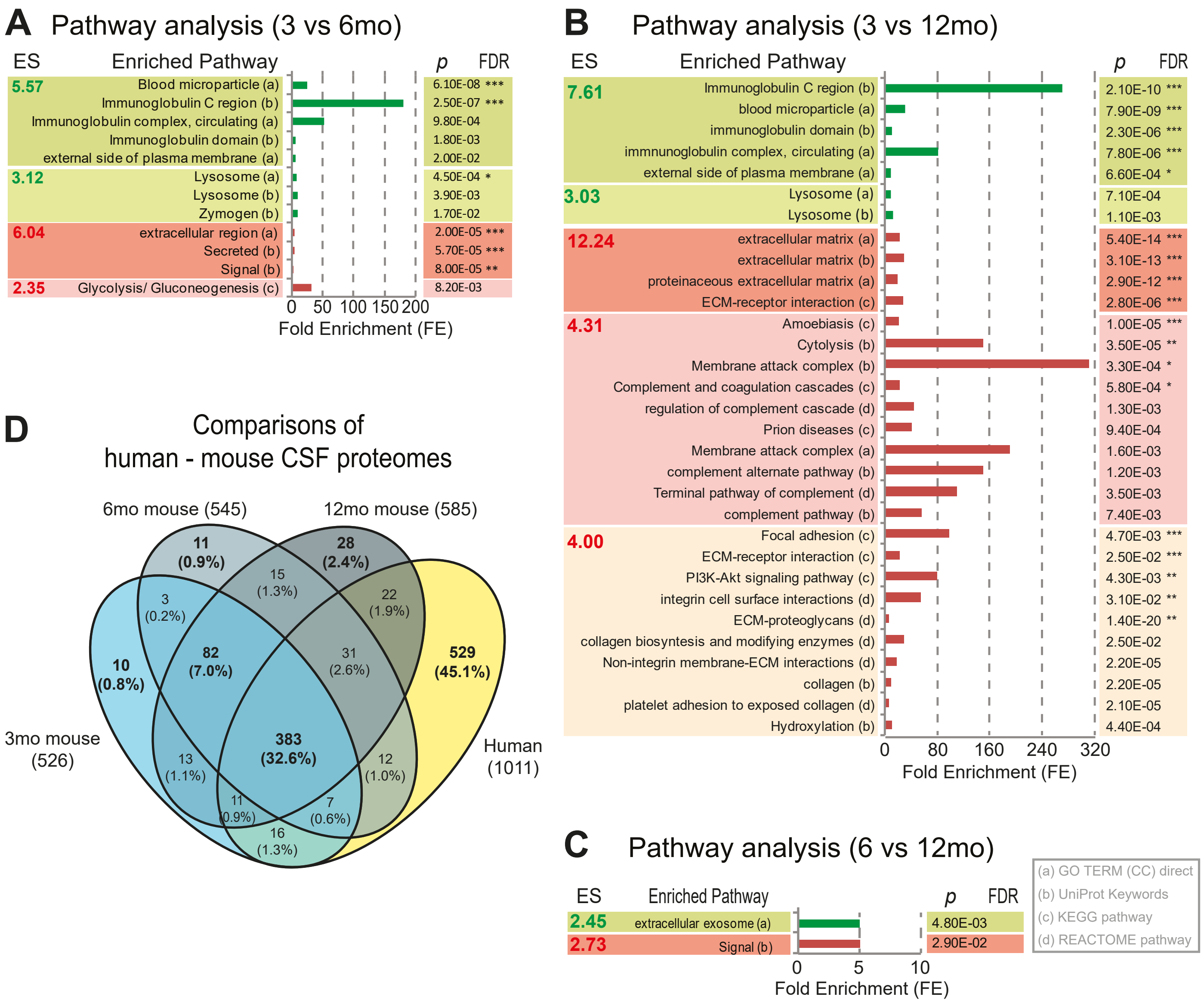


**Supplementary Figure 1.** Supplementary data to main Figure 2 for the analysis of single mouse CSF sampling. Pathway analysis of changed proteins between the A) 6 versus 3 months old, B) 12 versus 3 months-old and C) 12 versus 6 months-old mice, using the functional annotation clustering tool of DAVID-Database. The Enrichment Score (ES) ranks the overall importance (enrichment) of an annotation pathway in the proteome; green boxes contain the increased pathways, red boxes the decreased ones; the columns on the right indicate the corresponding p- (Benjamini correction) and FDR- (false discovery rate) significance values for each enriched (i.e. changed) pathway (*<0.05, **<0.01, ***<0.001); Fold Enrichment (FE) shows the magnitude of enrichment for each pathway (increased or decreased); the grey box next to (C) annotates via letters "a" to "d" the corresponding database for each of the enriched pathways. D) illustrative comparison (Venn diagram) of the common and different proteins between the 3 age-groups in mice and human CSF: 72.2% of the proteome remains common through aging from 3 to 12 months, the remaining 27.8% change dynamically.

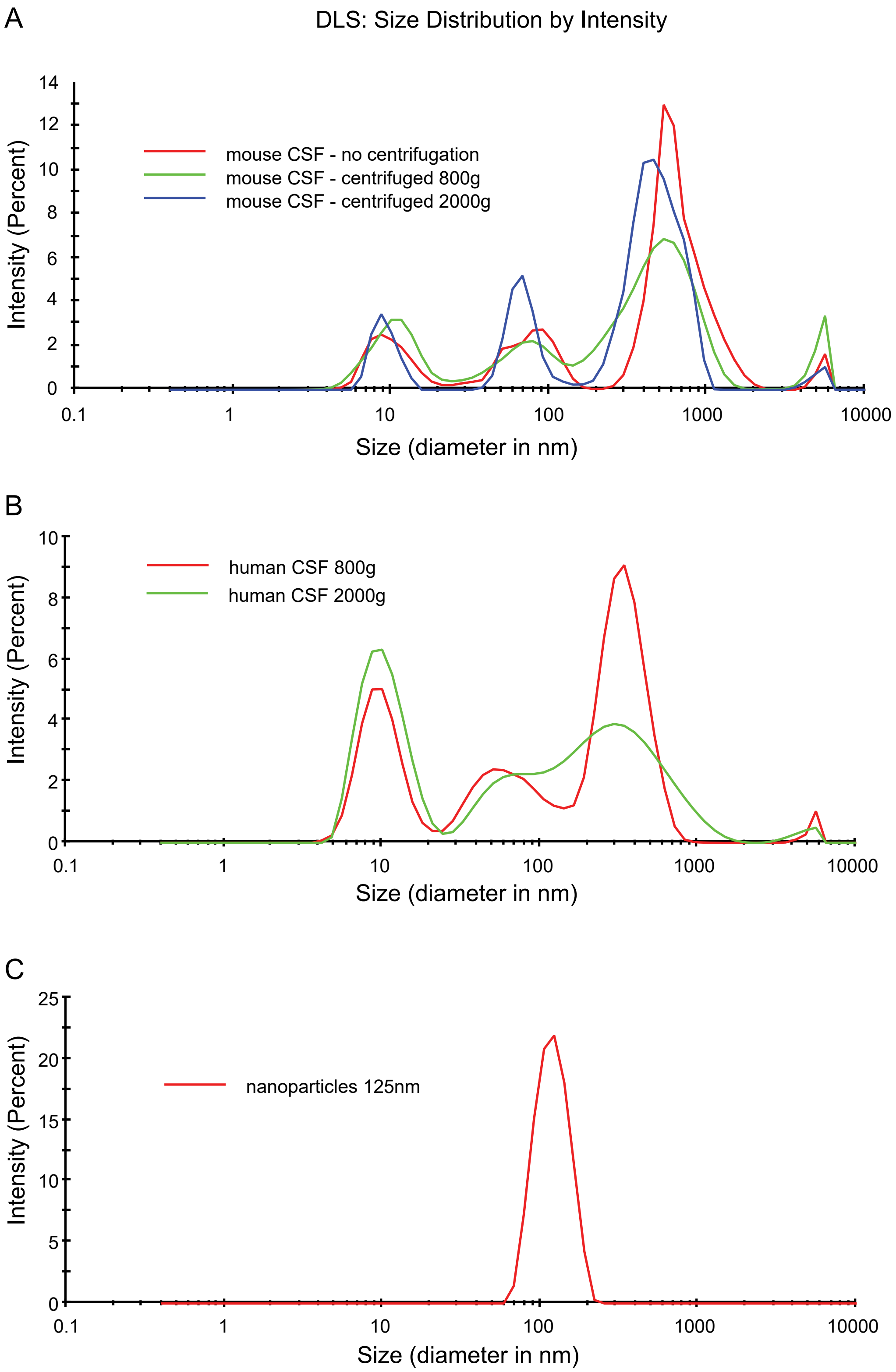


**Supplementary Figure 2.** Dynamic light scattering analysis of mouse and human CSF microparticles. Graphs show the size distribution of particles in mouse (A) or human CSF (B) under different centrifugation conditions. For assay quality control we use nanoparticles of a known size (C).

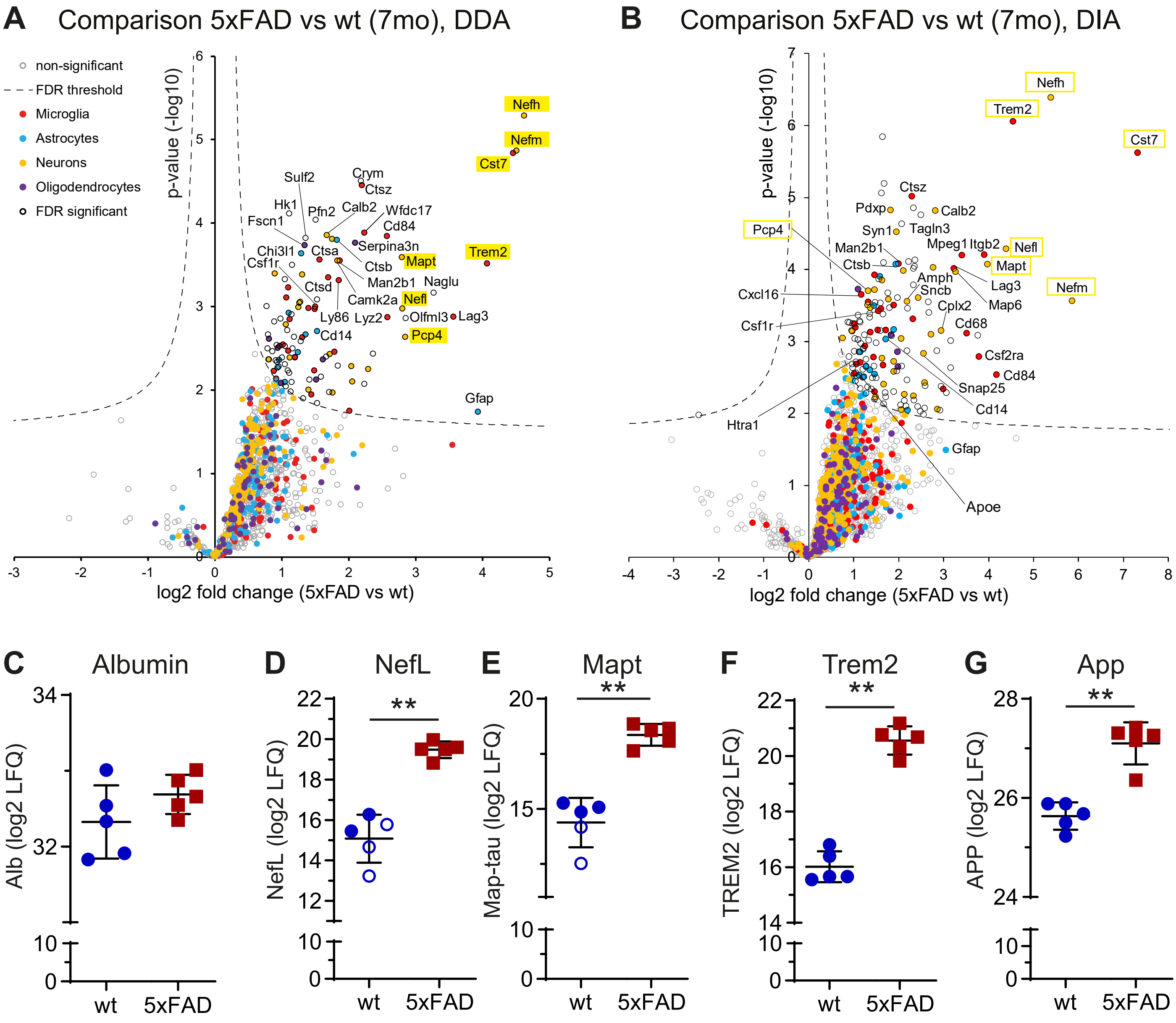


**Supplementary Figure 3.** A-B) Volcano plots of DDA (A) and DIA (B) analysis. The minus log10 transformed p-values of each protein are plotted against the log2 protein LFQ ratios. The dashed lines indicate a permutation-based FDR correction (p=0.05; s0=0.1). Proteins are categorized and labeled by color, based on their major cell type of origin (24). Selected and neurodegeneration-relevant proteins are highlighted in yellow boxes. LFQ quantifications for selected proteins of main Figure 5c, using the DIA analysis. C) Albumin CSF levels [log2(LFQ) values] did not differ between the 2 groups. Neurofilament light chain (NefL, D), MAPT (E), TREM2 (F) and APP (G) are significantly increased in 5xFAD mice compared to their corresponding littermates, verifying neurodegenerative and inflammatory activation in the CSF of the animals. ** p<0.01; line and whiskers show mean ±95% CI of each group. Imputed values are indicated as unfilled circles.
